# SpeSpeNet: An interactive and user-friendly tool to create and explore microbial correlation networks

**DOI:** 10.1101/2024.07.17.603889

**Authors:** A.L. van Eijnatten, L. van Zon, E. Manousou, M. Bikineeva, J. Wubs, W. van der Putten, E. Morriën, B.E. Dutilh, B.L. Snoek

**Affiliations:** Theoretical Biology and Bioinformatics, Science4Life, Utrecht University (UU), Padualaan 8, 3584 CH, Utrecht, the Netherlands; Institute for Biodiversity and Ecosystem Dynamics, Department of Ecosystem and Landscape Dynamics (IBED-ELD), University of Amsterdam, 1090 GE Amsterdam, the Netherlands; The Netherlands Institute of Ecology (NIOO); Institute of Biodiversity, Faculty of Biological Sciences, Cluster of Excellence Balance of the Microverse, Friedrich Schiller University, Jena, 07745, Germany

## Abstract

Correlation networks are commonly used to explore microbiome data. In these networks, nodes are taxa and edges represent correlations between their abundance patterns across samples. As clusters of correlating taxa (co-abundance clusters) often indicate a shared response to environmental drivers, network visualization contributes to system understanding. Currently, most tools for creating and visualizing co-abundance networks from microbiome data either require the researcher to have coding skills, or they are not user-friendly, with high time expenditure and limited customizability. Furthermore, existing tools lack focus on the relationship between environmental drivers and the structure of the microbiome, even though many edges in correlation networks can be understood through a shared relationship of two taxa with the environment. For these reasons we developed SpeSpeNet (Species-Species Network, https://tbb.bio.uu.nl/SpeSpeNet), a practical and user-friendly R-shiny tool to construct and visualize correlation networks from taxonomic abundance tables. The details of data preprocessing, network construction, and visualization are automated, require no programming ability for the web version, and are highly customizable, including associations with user-provided environmental data. Here, we present the details of SpeSpeNet and demonstrate its utility using three case studies.

## Introduction

The advent of high-throughput sequencing techniques and specifically amplicon-based sequencing has revolutionized microbial ecology (Semenov, 2021; Wensel et al., 2022). While research in this field used to be limited to culturable organisms, it is now possible to unravel the entire breadth of microbial diversity in any environment. Amplicon sequencing techniques have quickly become so accessible that data can now be generated at a faster pace than can be analyzed, while bioinformatic analyses often still require substantial programming skills (Hugerth & Andersson, 2017). Moreover, recent advances also allow shotgun metagenome data to be used for accurate taxonomic profiling (Hauptfeld et al., 2024).

Amplicon sequencing is based on the targeted amplification of marker sequences directly from the environment (Woese et al., 1985; Woese & Fox, 1977). The observed sequences are grouped into either Operational Taxonomic Units (OTUs) based on a percent similarity cutoff (Kozich et al., 2013), or Amplicon Sequencing Variants (ASVs) based on an error correction model that provides single nucleotide resolution (Callahan et al., 2017). The best-known marker sequence is the small subunit ribosomal RNA (Hadziavdic et al., 2014; Woese et al., 1985; Woese & Fox, 1977), suited for investigating bacterial and archaeal (16S) or eukaryotic (18S) communities. Another commonly used marker sequence is the internal transcribed spacer (ITS), suited for investigating the fungal community. Such marker sequences are highly conserved and can therefore be used to assign taxonomy by mapping the sequence of each OTU/ASV to a reference database such as Silva (Glöckner et al., 2017) or GreenGenes (DeSantis et al., 2006). The normalized marker sequence read counts can be used as a proxy for the relative abundance of the microbes in the environmental sample.

The relationships between microbes are often studied and visualized using network analysis (Dundore-Arias et al., 2023; Kishore et al., 2023; Ma et al., 2020; Morriën et al., 2017; Ramirez et al., 2018; Röttjers & Faust, 2018). Especially correlation networks are valuable tools to explore microbiome data. In such networks, nodes represent taxonomic lineages, either at the OTU/ASV level, or at a higher rank such as genus. Edges represent correlations between them, usually with a certain cutoff. While correlations between the relative abundance of microbial taxa should not be interpreted as direct interactions (such as cross-feeding, predation, competition, etc), they generally indicate a similar response to changes in the biotic or abiotic environment. Hence, clusters of co-occurring taxa (co-abundance clusters) often result from a shared response to one or more environmental drivers. Despite the generally high impact of environmental drivers on the structure of the microbiome, there are to our knowledge no network-visualization tools with adequate emphasis on the relationship between co-abundance clusters and environmental drivers.

Furthermore, current microbiome network tools are often inaccessible or impractical for many researchers. Microbiome networks may be constructed using general-purpose tools and R packages such as network (Butts, 2008), igraph (Csardi & Nepusz, 2006), Cytoscape (Shannon et al., 2003) or Gephi (Bastian et al., 2009). R-based packages for making networks specifically from microbiome data include NetCoMi (Peschel et al., 2021), microeco (Liu et al., 2021), and ggClusterNet (Wen et al., 2022). However, these still require programming ability on the part of the researcher, and it is often difficult to adjust the details and focus of the visualizations. The galaxy-based iNAP (Feng et al., 2022) does not require any programming skills but is inconvenient to work with due to the limitations of the galaxy platform and the lack of customization of the networks. Taken together, there is currently no user-friendly software to make correlation-networks from OTU/ASV tables and visualize environmental drivers of the microbiome.

To enable more researchers to make and customize networks, we introduce SpeSpeNet (Species-Species Network, https://tbb.bio.uu.nl/SpeSpeNet), an R-shiny based web-application that makes correlation-network visualizations directly from OTU/ASV tables. SpeSpeNet accelerates discovery and facilitates researchers by automating and customizing the data preprocessing, the construction of the network, and the network visualization. Using SpeSpeNet is intuitive and requires no programming ability. The taxa in the network can be colored by taxonomy, by relationship with user-defined environmental parameters, or by co-abundance clusters. SpeSpeNet also uses other visualization techniques, such as traditional taxonomy bar plots and boxplots showing the relationship between co-abundance clusters and environmental parameters. Here, we present the details of the SpeSpeNet tool and demonstrate its utility in microbial ecology with three case studies.

## Results

### Making correlation networks with SpeSpeNet

SpeSpeNet (Species-Species Network) is a fast, user-friendly, and comprehensive tool for visualizing and analyzing microbiome data using correlation networks. SpeSpeNet has two tabs that each visualize different aspects of the dataset: the network tab (**Figure S1**) and the summary tab (**Figure S2**). In the network tab, SpeSpeNet displays a correlation network where nodes are taxa and edges are correlations with a user-specified cutoff. We used a study on the microbial community in a bioremediation pipeline (Hauptfeld et al., 2022) (discussed in more detail in the next section) to showcase the networks made by SpeSpeNet. To explore the relationship between taxonomy and environmental drivers in the network, SpeSpeNet can color the nodes using four types of variables: 1) The taxonomy of the nodes at a chosen rank (e.g. order) (**Figure 1A**); 2) for a continuous environmental variable (e.g. O_2_), its correlation with the nodes (**Figure 1B**); 3) for a categorical environmental variable (e.g. location in the pipeline), the categories where nodes have the highest relative abundance (**Figure 1C);** and 4) the nodes’ co-abundance clusters, detected based on customizable settings (**Figure 1D**). The network is interactive: it shows the full taxonomy when the mouse is hovered over a node. Furthermore, SpeSpeNet can use categorical variables in the metadata to make a subnetwork based on a subset of the samples. This subnetwork feature allows the researcher to compare community structure and environmental drivers in contrasting environments.

**Figure 1:**
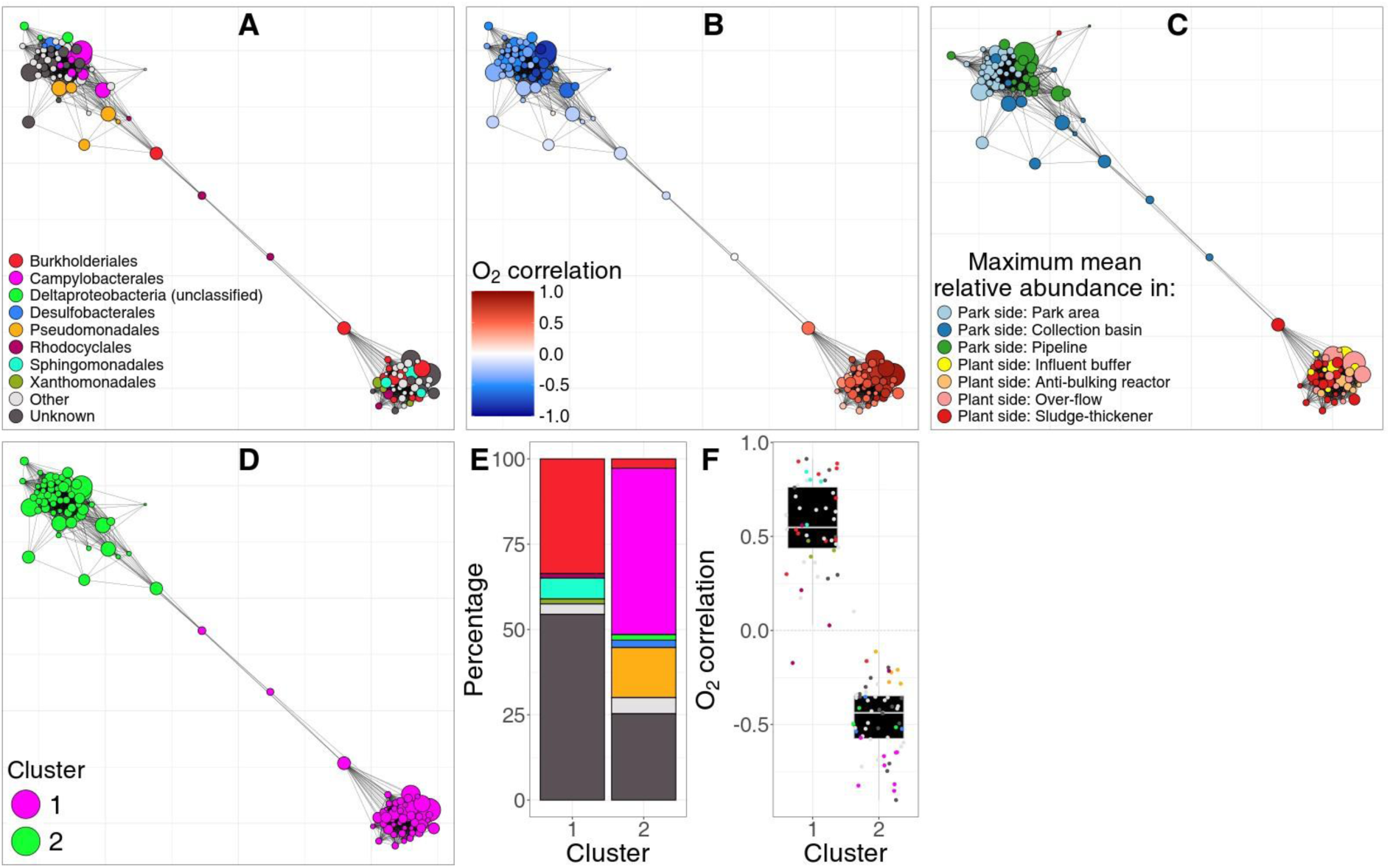
SpeSpeNet co-abundance network of the microbiome in a poly-contaminated city park and the corresponding water treatment plant. (Hauptfeld et al., 2022). Nodes are genera and edges are Spearman correlations > 0.63. **A)** Network colored by taxonomic annotation at the order rank. **B)** Network colored by Pearson correlation with O_2_ levels. The top-left cluster consists of genera that prefer lower O_2_ levels, and the bottom-right cluster corresponds to genera preferring higher oxygen levels. **C)** Network colored by the sampling site in the park/remediation pipeline in which genera were most abundant. The genera in the top-left cluster prefer the park side and the microbes in the bottom-right cluster prefer the plant-side. **D)** Network colored by k-means cluster assignment with k = 2. The k-means algorithm correctly groups genera into two co-abundance clusters that were already visually obvious. **E)** Barplot of taxonomic composition at order rank for the co-abundance clusters from D. Colors as in A. **F)** Distribution of correlation values with O_2_ for the taxa in the clusters from D. Co-abundance cluster 1 shows the microbial community in the park side, and cluster 2 in the plant side of the remediation pipeline. Colors as in A.

In the summary tab, SpeSpeNet offers an extended view of the clusters/categories in the network in terms of the taxonomic composition and the relationship to environmental drivers. This is achieved using two figures: 1) a barplot of the taxonomic composition of clusters or categories in the network (**Figure 1E**) and 2) a boxplot of the correlations of the nodes in each cluster or category with a selected environmental variable (**Figure 1F**).

SpeSpeNet provides many options to customize network generation in terms of the input format, network parameters, and aesthetic layout. Data can be input as .txt files or phyloseq objects (McMurdie & Holmes, 2013) (see **Supplementary text S1** for details on the required data format), or can be downloaded from the MGnify database (Richardson et al., 2023) into memory by SpeSpeNet. SpeSpeNet provides options to filter the taxa based on presence/absence and relative abundance thresholds. Additionally, various network options can be specified: 1) the type of correlation (Pearson, Spearman or Kendall) used to calculate the correlations between taxa; 2) the correlation cutoff that defines the edges in the network; and 3) the network clustering algorithm and parameters. Furthermore, SpeSpeNet lets the user specify the aesthetics of the network (**Figure S3**): width, transparency and arc of edges, the color of the background, and whether or not to plot isolated nodes (nodes without edges). A new network is automatically generated in only a few seconds when any of these options are changed.

Having discussed the functionalities and capabilities of SpeSpeNet, we now demonstrate its utility using three case studies. First, we used SpeSpeNet to investigate a dataset about the microbial community in a hydrocarbon-contaminated park and the associated treatment plant (Hauptfeld et al., 2022). Second, we applied SpeSpeNet to a dataset of marine microbiome samples from the north-western Mediterranean (Blanes Bay, Spain) (Deutschmann et al., 2023) to investigate phytoplankton blooms. Finally, we used SpeSpeNet to analyze a dataset about the effect of different organic amendments on the soil microbiome and microbial functional gene abundance (Brenzinger et al., 2021).

#### Case study 1: Network of the microbial community in a poly-contaminated park and the associated treatment plant

We investigated the microbial community in a hydrocarbon-contaminated park and the associated treatment plant. The samples came from seven different stages of the bioremediation process. These samples could be broadly subdivided into anaerobic soil samples, representing the “park-side” of the system (stages 1-3) and samples from the aerated water treatment “plant-side” (stages 4-7). The network made by SpeSpeNet clearly recapitulated two types of microbiomes as separate co-abundance clusters: one dominated by taxa of the order *Burkholderiales*, the other dominated by taxa of the order *Campylobacterales* (**Figure 1A**).

We used SpeSpeNet to investigate potential environmental drivers of the two co-abundance clusters visible in the network. Coloring the taxa based on their Pearson correlation with oxygen (O_2_) showed that the two clusters in the network are linked to O_2_ levels (**Figure 1B**). This result aligned with the conclusions by Hauptfeld *et al.:* In the water treatment plant, the added oxygen facilitated the change from an anaerobic to an aerobic microbial community. The top-left co-abundance cluster in the network consisted exclusively of microbes that correlated negatively with O_2_ levels (anaerobic community), whereas the bottom-right co-abundance cluster consisted of microbes that correlated positively with O_2_ levels (aerobic community). SpeSpeNet further showed that the taxa in the anaerobic community were most abundant in the park-side stages, whereas the taxa in the aerobic community were most abundant in the treatment plant-side stages (**Figure 1C**).

Within SpeSpeNet, the k-means clustering could distinguish between the two different communities (**Figure 1D**). In this way, SpeSpeNet visualized the taxonomic composition of the two communities (**Figure 1E**). The barplots showed that more than half of the park-side community consisted of taxa belonging to the orders *Campylobacterales* and *Pseudomonadales*, while most of the plant-side community consisted of taxa belonging to the orders *Burkholderiales* and *Sphingomonadales*. Finally, SpeSpeNet visualized the distribution of Pearson correlation with O_2_ levels in each community in a boxplot (**Figure 1F**), confirming their inverse correlation with O_2_ levels.

#### Case study 2: Phytoplankton-blooms in the north-western Mediterranean

As a second case study, we used a dataset of marine microbiome samples from the north-western Mediterranean (Blanes Bay, Spain) (Deutschmann et al., 2023). In this study, the surface water was sampled monthly for ten consecutive years and the microbial community composition was determined using the 16S and 18S marker genes. The 16S and 18S tables were merged after rarefaction to create a single abundance table, needed to calculate correlations between all taxa. Several environmental factors were also measured, including temperature and total chlorophyll a. Chlorophyll a is considered a proxy for the phytoplankton biomass (Jakobsen & Markager, 2016) and it peaks during phytoplankton blooms (Gutiérrez-Rodríguez et al., 2010). We studied the taxa involved in phytoplankton blooms in Blanes Bay by correlating taxa relative abundance with chlorophyll a.

First, we sanity checked the data using SpeSpeNet by making barplots of the prokaryotic and eukaryotic microbiome in each month and investigating if the observed patterns match known seasonal patterns from the area (**Figure 2A-B**). The barplots show several patterns that are in agreement with previous literature, such as the increase in *Synechococcales* between July and Oktober (Gasol, 2016) (**Figure 2A**) and the strong seasonality of the *Mamiellales* (Giner et al., 2019) (**Figure 2B**). Next, we used SpeSpeNet to make an inter-kingdom correlation network using the merged 16s and 18s abundance tables (**Figure 2C**). Because the samples varied in time but not in space, the correlations showed a cyclical structure that corresponded to the changing of environmental conditions, such as temperature and nutrient levels, during the passing of the seasons (**Figure 2D-E, Figure S4**).

**Figure 2:**
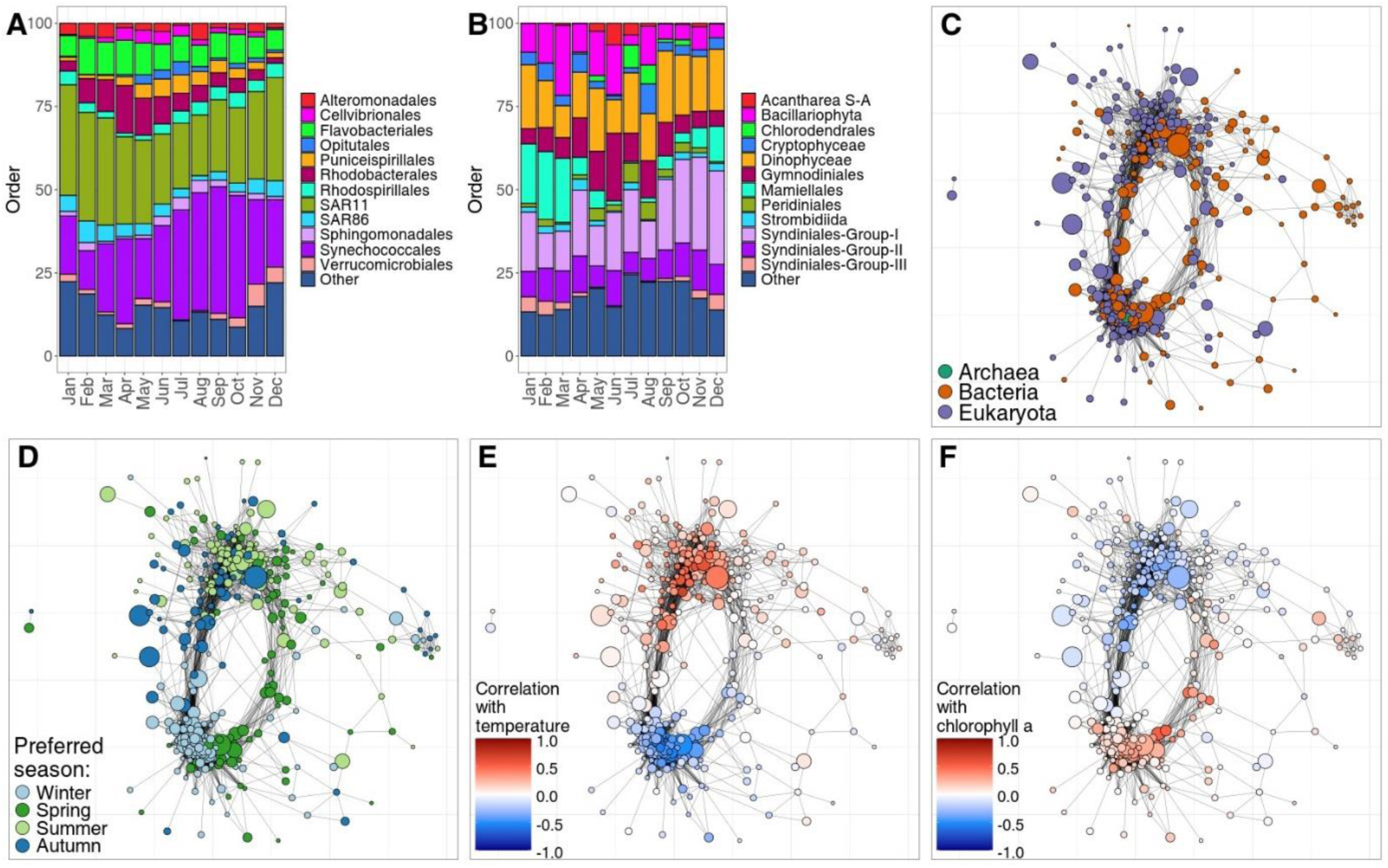
Inter-kingdom associations in a 10-year longitudinal study in the north-western mediterranean (Deutschmann et al., 2023). Abundance table made by merging 16S and 18S table. In the networks, nodes are genera and edges represent Spearman correlations > 0.4. **A)** Taxonomic composition of the prokaryotic community per month at order rank. **B)** Taxonomic composition of the Eukaryotic community per month at order rank. **C)** Co-abundance network colored by the kingdom of each genus. **D)** Network colored by the season during which taxa have the highest mean relative abundance. **E)** Network colored by the correlation between each genus and surface water temperature. **F)** Network colored by the correlation between each genus and chlorophyll a levels.

Next, we colored the network by the correlation with chlorophyll a (**Figure 2F**). Mousing over nodes with high correlations in SpeSpeNet showed that some belonged to the families *Roseobacteraceae* (Liang et al., 2021) and *Flavobacteraceae*. These two families have been identified as the two main bacterial groups associated with phytoplankton blooms (Buchan et al., 2014). In the case of *Roseobacteraceae*, the association with phytoplankton blooms results from cross-feeding interactions with micro-algae involved in phytoplankton blooms (Suleiman et al., 2016; Zecher et al., 2020). To quantify the associations between taxa relative abundance and chlorophyll a levels, we used the spring and summer samples to calculate Pearson correlations. Out of the six prokaryotic genera with the highest correlations with chlorophyll a, five belonged to *Roseobacteraceae* (*Amilybacter*, *Loktanella* and *Planktomarina*) and *Flavobacteraceae* (*Aurantivirga* and *Winogradskyella*).

The top three eukaryotic genera with the highest correlations with chlorophyll a were diatoms (*Pseudo-nitzschia*, *Thalassiosira* and unknown *Coscinodiscophyceae*). Members of *Roseobacteraceae* have been found in the microbiome of various diatoms including *Pseudo-nitzschia* (Koester et al., 2022) and *Thalassiosira* (Tran et al., 2023). Therefore, we used SpeSpeNet to identify *Roseobacteraceae* genera that strongly correlated with the relative abundance of these two diatom genera (Spearman correlation > 0.6). We found that *Planktomarina* and *Amylibacter* strongly correlated with *Pseudo-nitzschia* (ρ = 0.68 and ρ = 0.62, respectively). Potentially, bacteria of *Planktomarina* and/or *Amylibacter* could be involved in metabolic interactions with *Pseudo-nitzschia* diatoms. This case study about phytoplankton blooms shows that SpeSpeNet captures cyclic seasonal dynamics and can be used to form hypotheses about interkingdom interactions.

#### Case study 3: Influence of organic amendments on the soil microbiome and greenhouse gas emissions

For the third case study, we use a dataset about the effect of different organic amendments on the soil microbiome and microbial functional gene abundance (Brenzinger et al., 2021). Six different amendments were applied to sand and clay soils: Residue derived from cover crops harvested in Februari (Ccfeb) or November (Ccnov), a mixture of digestate and compost (DC), a mixture of sewage sludge and compost (SC), mineral fertilizer (MF), and a control (control65). Half of the plots were planted with *Triticum aestivum* (common wheat). We used SpeSpeNet to make a network of the soil microbiome and colored by the treatment in which taxa have the highest mean relative abundance. We noticed two main clusters in the network, both with a mixture of microbes favoring different amendments (**Figure 4A**). Using the soil type to color the network showed that the two clusters corresponded to microbes favoring sand or clay soils (**Figure 4B**). To reveal the effect of the amendments on the microbiome through the confounding effect of soil type, we used SpeSpeNet to automatically make a subnetwork using only the samples annotated as clay.

**Figure 4:**
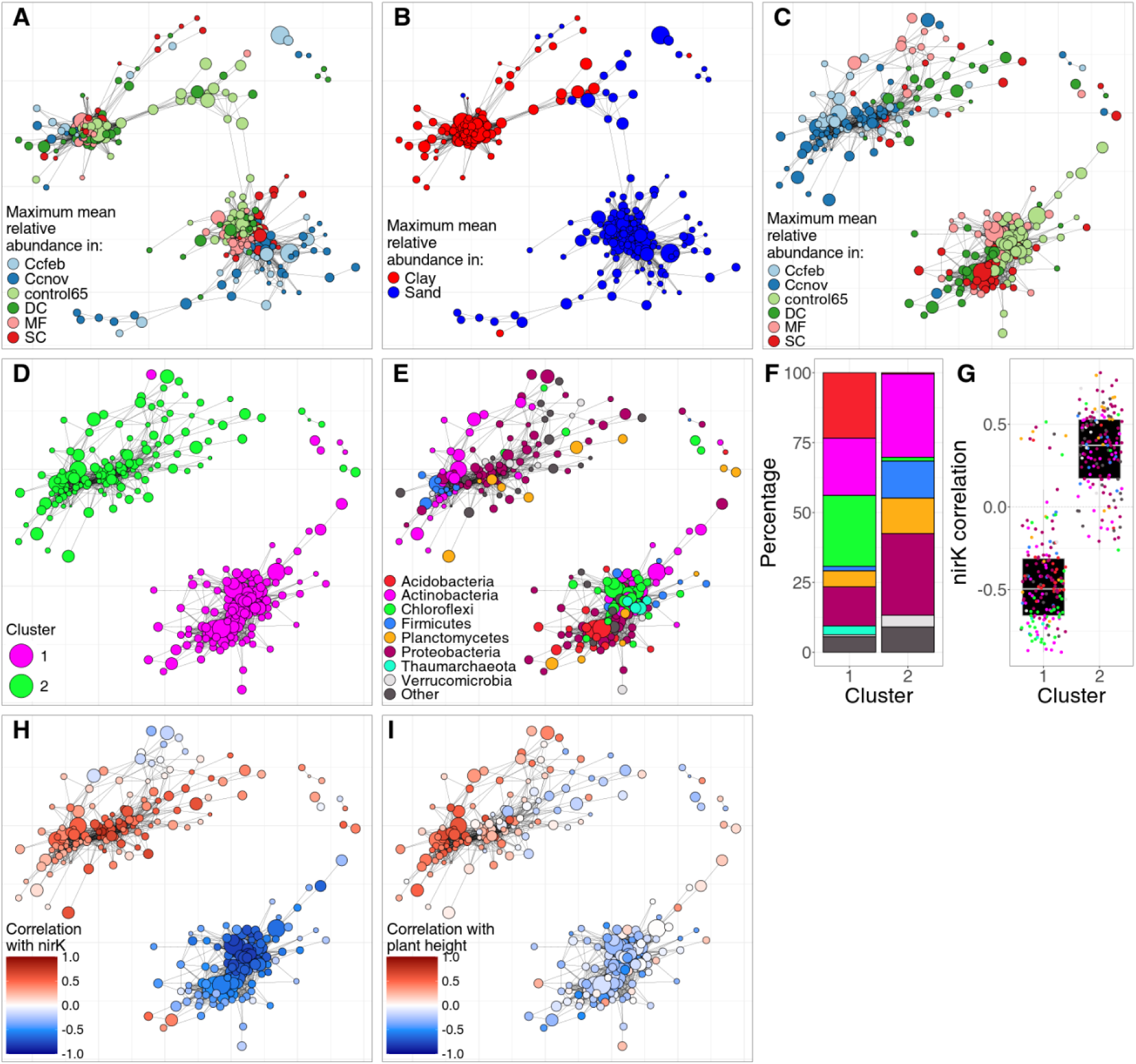
Effect of amendments on the microbiome in a mesocosm experiment (Brenzinger et al., 2021). **A)** Network using all samples, colored on the amendment condition in which taxa have the highest relative abundance. Nodes are genera and the Spearman correlation cutoff is 0.72. **B)** Same network as in A colored on the soil type in which genera have the highest relative abundance. **C)** Subnetwork using only clay samples, colored by the amendment condition in which taxa have the highest relative abundance. Nodes are genera and the Spearman correlation cutoff is 0.58. **D)** Subnetwork colored by k-means clustering with k=2. **E)** Subnetwork colored by taxonomy at phylum rank. **F)** Barplot of the taxonomic composition at phylum rank of the two co-abundance clusters shown in D. **G)** Distribution of the correlation with the qPCR abundance of the *nirK* genes for the genera in the co-abundance clusters shown in D. **H)** Subnetwork colored by correlation of relative abundance of genera with the qPCR abundance of the *nirK* gene (involved in NO_2_ reduction). **I)** Subnetwork colored by correlation of relative abundance of genera with the plant height after the experiment.

We created a subnetwork to investigate the effect of amendments on the microbiome in clay soils (**Figure 4C**). SpeSpeNet shows two co-abundance clusters: One cluster contains all taxa that favor the control65 (control cluster) and the other cluster contains almost all taxa that favor the Ccfeb and Ccnov amendments (cover crop cluster). Taxa that favor the SC, DC or MF amendments are more spread between the co-abundance clusters. This shows that especially the cover crop-based amendments favor distinct taxa compared to unamended soils. To further investigate the differences in microbial community between the co-abundance clusters, we detected the co-abundance groups using the k-means algorithm (**Figure 4D**) and plotted the respective taxonomy (**Figure 4E**, **Figure 4F**). The taxonomy differs strikingly between the co-abundance clusters, even at phylum rank, with the cover crop cluster 2 being enriched for *Proteobacteria* and *Firmicutes* but depleted for *Acidobacteria* and *Chloroflexi*.

The dataset also contained qPCR abundances of various functional marker genes. We wondered whether the different taxonomic compositions of the two co-abundance clusters translated to different community functions. Therefore, we investigated whether the taxa in the two co-abundance clusters had different correlations with the qPCR abundances of functional genes related to nitrogen cycling. Various nitrogen cycling genes such as *NirK* (NO_2_ reduction) (**Figure 4G**, **Figure 4H**), *NifH* (nitrogen fixation) (**Figure S5A**), and *NOSZII* (N_2_O reduction) (**Figure S5B**) are strongly associated with the two clusters in the network. This could indicate that the co-abundance clusters in the network are largely driven by the impact of amendments on the nitrogen availability in the soil. Interestingly, a subset of the microbes in the cover crop cluster, which positively correlates with increased nitrogen cycling genes, have strong positive correlations with plant height (**Figure 4I)**. Mousing over the network in SpeSpeNet shows that these include known plant beneficial nitrogen-fixing genera such as *Lysobacter* (**Figure S6A**) and *Achromobacter* (**Figure S6B**). In this case study, we show how SpeSpeNet can be applied to rich datasets with different data types to quickly form hypothesis about the interplay between the environment, microbiome, microbial function, and other biota.

## Discussion

Here, we present SpeSpeNet, an interactive network visualization tool that facilitates visualization and investigation of co-abundance networks by streamlining the network construction process from OTU/ASV count tables. With three different use cases, we showed how these networks allow discovery of meaningful microbiome-environment associations. The networks are highly customizable in terms of the network parameters and aesthetics. Furthermore, network construction only takes a few seconds, even if hundreds of samples are used as input. All of this can be done at the click of a button and requires no programming skill on the part of the user.

Currently, several alternative methods to SpeSpeNet also construct networks from microbiome data. For example, the R-package NetCoMi (Peschel et al., 2021) handles data filtering, normalization, and network construction. However, this package still requires the researcher to have some technical ability in programming. Furthermore, it does not offer the same focus on the relationship between environmental drivers and the microbiome, and the network visualizations are not of the same aesthetic quality as those generated by SpeSpeNet. Other R-based approaches such as microeco (Liu et al., 2021) and ggClusterNet (Wen et al., 2022) have similar limitations. As far as we know, the only other approach that requires no programming from researchers is the galaxy-based iNAP (Feng et al., 2022). However, this approach also lacks focus on the relationship between the microbiome and environmental drivers and produces less aesthetic visualizations compared to SpeSpeNet. Furthermore, the galaxy pipeline can be hard to troubleshoot if an error is given at any stage. Therefore, the user-friendliness, the focus on environmental drivers, and the quality of the visualizations make SpeSpeNet a valuable addition to the current microbiome network analysis approaches.

SpeSpeNet visualizes associations between the microbiome and the environment by correlating taxa with environmental variables. Moreover, SpeSpeNet shows the distribution of these correlations for the identified co-abundance clusters, pinpointing potential drivers of the microbiome. We stress that such patterns, while they can be insightful, always need to be interpreted with caution. After all, environmental variables are often highly (anti-)correlated with each other, such that even if only one variable is driving the microbiome, many different variables will show a similar, visually convincing pattern (**Figure S7**). Furthermore, correlations cannot inform on the direction of the effect, and therefore cannot show whether the microbiome is driving the environment, or the environment is driving the microbiome. Thus, SpeSpeNet should be used to explore and generate hypotheses rather than conclusions. These hypotheses can be further investigated using appropriate techniques and statistics, and inform or inspire targeted follow-up experiments.

Edges in the networks generated by SpeSpeNet should not be overinterpreted for two major reasons. Firstly, the correlation-cutoff defining an edge is chosen arbitrarily. Two taxa may have correlations that are barely higher than the cutoff, whereas two other taxa may have correlations that are just lower. The network will show an edge between the first two taxa but not the second two, despite the correlations being almost equal (although in either case they will likely end up in the same co-abundance cluster through correlations with other taxa). Thus, the discrete nature of the network does not reflect the continuous nature of the correlation matrix. Secondly, microbiome data is compositional in nature (Gloor et al., 2017). Read counts need to be normalized for variation in sampling depth, because the sampling depth is a purely technical signal. SpeSpeNet performs this normalization by scaling the read counts with the total sum of the reads in the sample. The result is that the sum of the reads in the sample is constrained to a fixed value. This compositionality has important consequences for data interpretation, because if one taxon increases in abundance, all others must necessarily decrease. This may induce spurious correlations. The effect to which the correlation network is impacted by compositionality will depend on the dataset. Datasets with high alpha-diversity (e.g. soil) are likely less impacted by compositionality than less diverse datasets (e.g. human skin). Furthermore, datasets that combine samples from different biomes that vary in total microbial abundance, are more likely to be impacted by compositionality than datasets where all samples are from a similar system with comparable microbial density.

Several “compositionally aware” normalization methods have been developed (Ha et al., 2020; Jiang et al., 2020; Kurtz et al., 2015; Quinn et al., 2017; Tackmann et al., 2019; Watts et al., 2019; Yoon et al., 2019) that attempt to correct for the spurious correlations induced by compositionality. However, different compositionally aware methods result in very different networks with little overlap in terms of edges (Kishore et al., 2023). Therefore, we implemented total sum scaling in SpeSpeNet because this has a clear interpretation and, to our opinion, captures the general patterns in terms of the interaction between the microbiome and environmental drivers well. Future versions of SpeSpeNet may include options for other normalization methods.

Researchers should exercise caution when interpreting edges ecologically. Edges indicate, per definition, that taxa correlate in their relative abundance. This rarely implies a direct interaction in the ecological sense, such as predation or cross-feeding (Barberán et al., 2012). Instead, taxa often simply respond in a similar way to external drivers, causing their relative abundances to change in similar directions, e.g. in response to a fluctuating environment. Alternatively, edges could result from a shared response to changes in community composition and its functional potential. Either scenario could cause a correlation between the relative abundance of two taxa without requiring a direct ecological interaction between them. However, elucidating what microbes respond similarly to environmental changes is highly relevant for topics such as the effect of climate change on the soil microbiome or the impact of a diet/drug on the gut-microbiome.

SpeSpeNet uses k-means on the correlation matrix to identify communities. This comes with several caveats, because the k-means algorithm is not explicitly designed for community detection. First, the k-means algorithm assigns all taxa to one of the clusters, regardless of whether a taxon has a clear co-occurrence pattern with any of the other taxa. Secondly, k-means assumes spherical clusters and cannot accurately detect communities with a more complex configuration in correlation-space. Care also needs to be taken when assigning a biological interpretation to the k-means clusters. For example, a user might interpret a k-means cluster as a group of microbes being driven by similar environmental drivers based on the boxplot in the summary tab. However, the k-means clusters can include outlier taxa that correlate differently with environmental data than others (as in e.g. **Figure 4G** cluster 1). If these outlier taxa happen to have a very high relative abundance, they will have a strong impact on the taxonomic composition of the k-means clusters (as shown in the barplots in the summary tab). In such cases, the user must take care not to interpret this change in taxonomic composition as being caused by or responsible for the environmental pattern. We emphasize that SpeSpeNet is a tool for ecological data visualization and hypothesis generation.

In the future, SpeSpeNet could be extended in several ways. Firstly, we want to provide the user with different options to normalize their data using methods that take compositionality into account such as CLR and ILR transformations. Secondly, we would like to implement more advanced network inference methods such as SparCC (Kishore et al., 2023) and EleMi (Chen & Bucur, 2024). Thirdly, we want to implement different subcommunity detection algorithms as alternatives to k-means (e.g. the Leiden algorithm (Traag et al., 2019)). Finally, we want to provide an option to the user to input functional assignments of the OTUs/ASVs (such as the FUNGuild (Nguyen et al., 2016) output for ITS sequences). In its current state, SpeSpeNet focuses on the interaction between the microbiome and the environment. However, the three-way interaction between the environment, the microbiome, and microbial functions would be even more informative, and could reveal additional relevant biology.

In summary, SpeSpeNet speeds-up and facilitates discovery in microbial ecology by enabling researchers to interactively explore their datasets through network visualization and clustering. SpeSpeNet puts particular emphasis on the detailed exploration of potential environmental drivers of microbial community composition as a whole or on the level of co-abundance clusters. For these reasons we believe that SpeSpeNet will be a valuable addition to the field of microbial ecology.

### Methods

#### Dependencies

SpeSpeNet is an Rshiny tool, developed with the shiny package (Chang W, 2023). The layout of the app is generated using the “cyborg” option from the shinythemes package (Winston Chang, 2021). Network visualizations are made with the igraph package (Csardi & Nepusz, 2006), using the ggraph package (Thomas Lin Pedersen, 2024a) for programmatic convenience. For other plots SpeSpeNet uses the ggplot2 package (Martin et al., 2022), supplemented by the ggdark package (Neal Grantham, 2018). SpeSpeNet uses the MGnifyR package (Tuomas Borman, n.d.) to download public data from the MGnify database into SpeSpeNet. The network and other plots are made interactive with the ggiraph package (David Gohel and Panagiotis Skintzos, 2023) and the plotly package (Plotly Technologies Inc., 2015) respectively. The code makes use of the tidyverse (Wickham et al., 2019) and tidygraph (Thomas Lin Pedersen, 2024b) packages for convenience.

#### Data format and upload

Data can be input as three separate tab delimited .txt files that contain the OTU/ASV table, the taxonomy table, and the environmental table or as a phyloseq object (.rds file). For the details on the data format SpeSpeNet requires see the manual (**Supplementary text S1**). SpeSpeNet can also download studies from the MGnify database if provided with a MGnify study accession number.

The user can choose to aggregate the OTUs/ASVs to genus rank during upload. In this case, the relative abundances of all taxa belonging to the same genus are summed and the nodes in the network will be genera instead of OTUs/ASVs. Taxa that cannot be determined at the genus rank are aggregated to the deepest known taxonomic rank. As an example, all taxa that can only be determined to the order *Burkholderiales* will be a single node in the network, possibly containing multiple genera or families.

#### Data processing

After the data is uploaded the read counts per OTU/ASV are normalized to relative abundances as a percentage of the total reads in the sample. Then, the taxa are filtered using two user-specified thresholds: 1) The minimum number of samples a taxon needs to occur in (have non-zero read counts) and 2) the minimum % of total reads a taxon needs to have in at least one of the samples. Only OTUs/genera that satisfy both thresholds are included in the network.

#### Network construction and visualization

The filtered matrix with relative abundances is used to calculate the correlation matrix with dimensions taxa by taxa. The default correlation method is Spearman, but SpeSpeNet also allows the user to choose Pearson or Kendall correlations. We recommend a rank-based correlation as these could be less sensitive to the compositional nature of the data.

SpeSpeNet then constructs a graph from the correlation matrix using the function graph.adjacency() from the igraph package. All edges between taxa with a correlation lower than the specified cutoff are deleted using the function delete.edges(). Next, the network is plotted using the ggraph() function from the ggraph package. The size of the nodes in the network are proportional to the square root of the mean relative abundance over all samples of the corresponding taxa. The user can choose whether to plot isolated nodes (nodes without edges). However, these nodes are still used in the k-means clustering. The user can also set the curvature, width, and transparency of the edges in the network.

SpeSpeNet can make a subnetwork using categorical variables in the metadata. After the user selects a categorical variable, SpeSpeNet lists all the categories in this variable that correspond to at least eight samples. After the user selects a category, SpeSpeNet will calculate the taxa by taxa correlation matrix and construct a network using only samples from that category.

The network can be colored on 1) k-means cluster assignment 2) relationship with environmental variable and 3) taxonomy of the nodes. In the first case, the k-means algorithm is run on the taxa by taxa correlation matrix with a user-specified K (number of clusters). Cluster assigned is then used to color the network. In the second case, the environmental variable can either be numeric or categoric. For a numeric environmental variable, the Pearson correlation between the relative abundance and the environmental variable is calculated for each taxon and used to color the network. NA values are discarded when calculating the Pearson correlations. For a categorical environmental variable, SpeSpeNet calculates the category in which each node has the highest mean relative abundance. This is then used to color the network. Finally, in the third case a user-specified taxonomic rank is used to color the nodes of the network. The fifteen clades with the highest relative abundance are shown, and all the others are labelled as “Other” to constrain the size of the legend and the number of colors in the network.

#### Summary tab

The summary tab shows two plots. The first is a barplot of the taxonomic composition at a chosen taxonomic rank. If the network is colored on a categorical environmental variable, the barplot will show the taxonomic composition per category. If the network is colored on k-means cluster, the barplot shows the taxonomic composition of each cluster separately. In this case, the user can choose whether they want to show the mean relative abundance as a percentage of the cluster or as a percentage of all reads. Finally, if the network is colored by taxonomy or a continuous environmental variable in the network tab, the barplot shows the mean relative abundance per clade over all samples.

The second plot in the summary tab is a boxplot of the Pearson correlations between the relative abundances of the taxa in the network with a continuous environmental variable of choice. If the network is colored on k-means, the boxplot is faceted on cluster.

## Data availability

SpeSpeNet is available as a webtool from https://tbb.bio.uu.nl/SpeSpeNet. The code for SpeSpeNet (which can be used to launch SpeSpeNet offline) and the code to make the figures in the manuscript are on https://github.com/SnoekLab/SpeSpeNet. The github also contains the data of the three case studies presented in this document as .txt files, formatted to be SpeSpeNet compatible. The SpeSpeNet manual can be found on the github and as Supplemental text S1.

## Supporting information

Supplemental text S1: SpeSpeNet manual

## Acknowledgements

This publication is part of the SoilProS project (funded by TTW Perspective Program P20-45 of the Dutch Research Council (NWO)). BED was supported by the European Research Council (ERC) Consolidator grant 865694: DiversiPHI, the Deutsche Forschungsgemeinschaft (DFG, German Research Foundation) under Germany’s Excellence Strategy – EXC 2051 – Project-ID 390713860, and the Alexander von Humboldt Foundation in the context of an Alexander von Humboldt-Professorship founded by German Federal Ministry of Education and Research.

## Author contributions

BLS conceived the idea with input from E. Morriën, JW, WvdP and BED. AlvE, BLS and BED wrote the manuscript with input from all co-authors. ALvE did the coding and design with help from LvZ and BLS. ALvE, E. Manousou and MB did bug and interaction testing. E. Manousou and MB helped with the case studies. E. Manousou wrote the manual with help from ALvE.

## Supplementary Materials

### Supplementary figures

**Figure S1:**
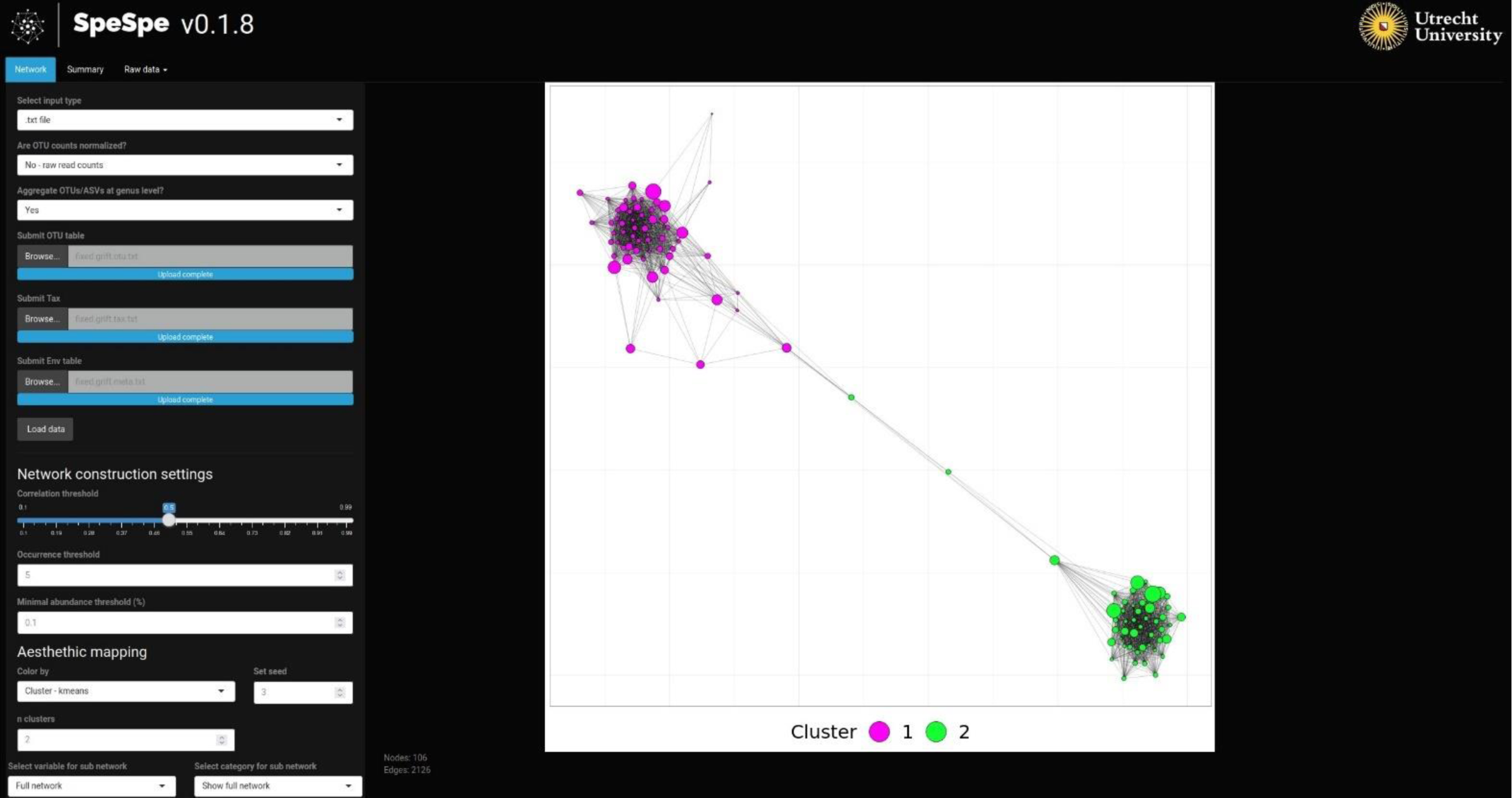
Screenshot of the network tab of the SpeSpeNet web application. Same data as in case study 1 (Hauptfeld et al., 2022). The network is colored on a k-means clustering with k = 2. On the left are various buttons and sliders that can be used to change the details of the data upload, data filtering and network construction. Some options are not shown as they did not fit on the screen. For zoomed in images of all the customization options see the SpeSpeNet manual (**Supplemental text S1** or on the github). The network generated after clicking “Load data” is shown on the right (all parameters are left on default except for the random seed and the number of clusters).

**Figure S2:**
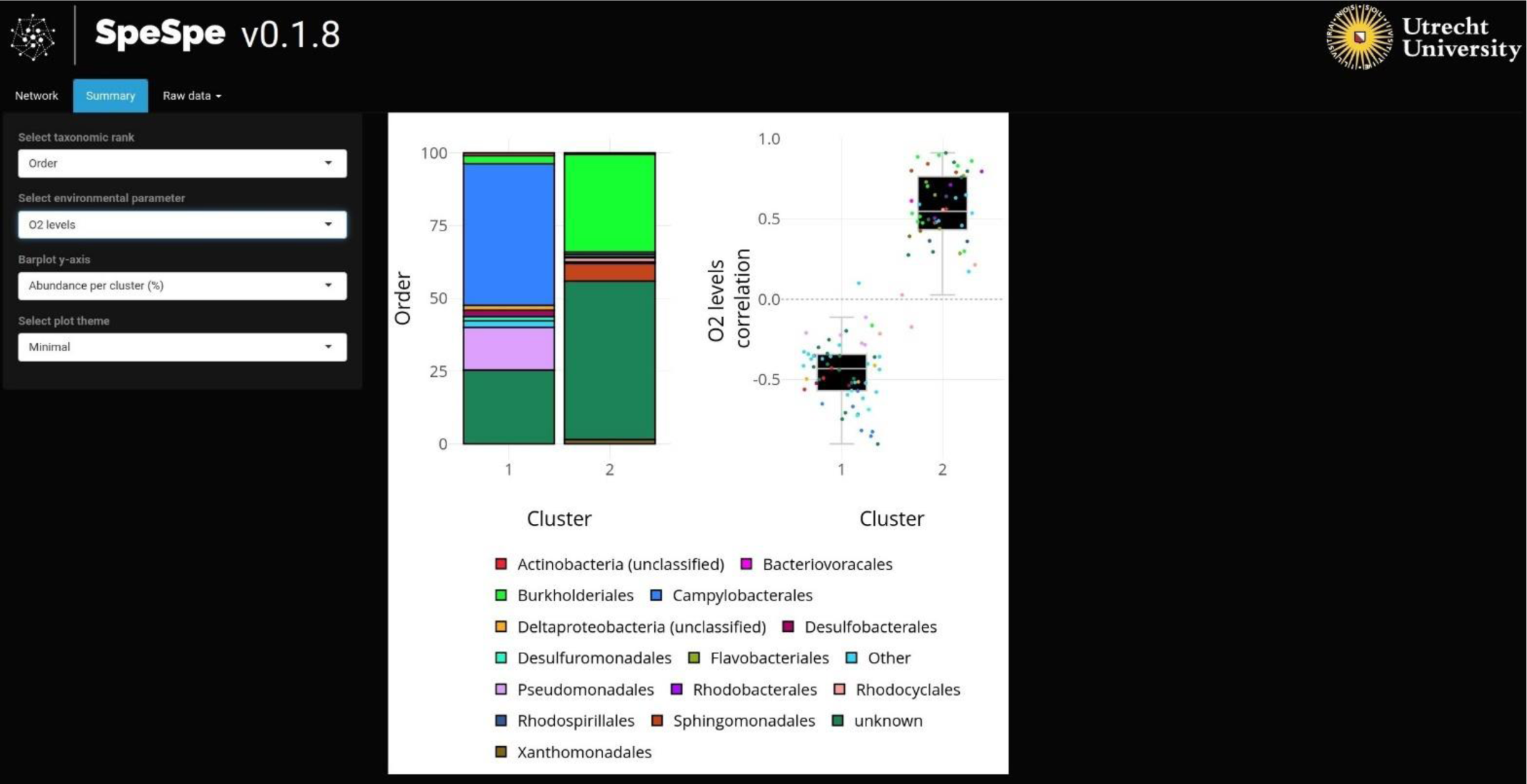
Screenshot of the summary tab of the SpeSpeNet web application. The data is the same as in case study 1 (Hauptfeld et al., 2022). On the left are options to change the taxonomic rank used for the barplot, the environmental variable for the correlations in the boxplot, the layout of the barplot, or the background theme of the figures. On the right is a barplot of the taxonomic composition at order rank and a boxplot with the correlation with O_2_ for the clusters shown in Figure S1.

**Figure S3:**
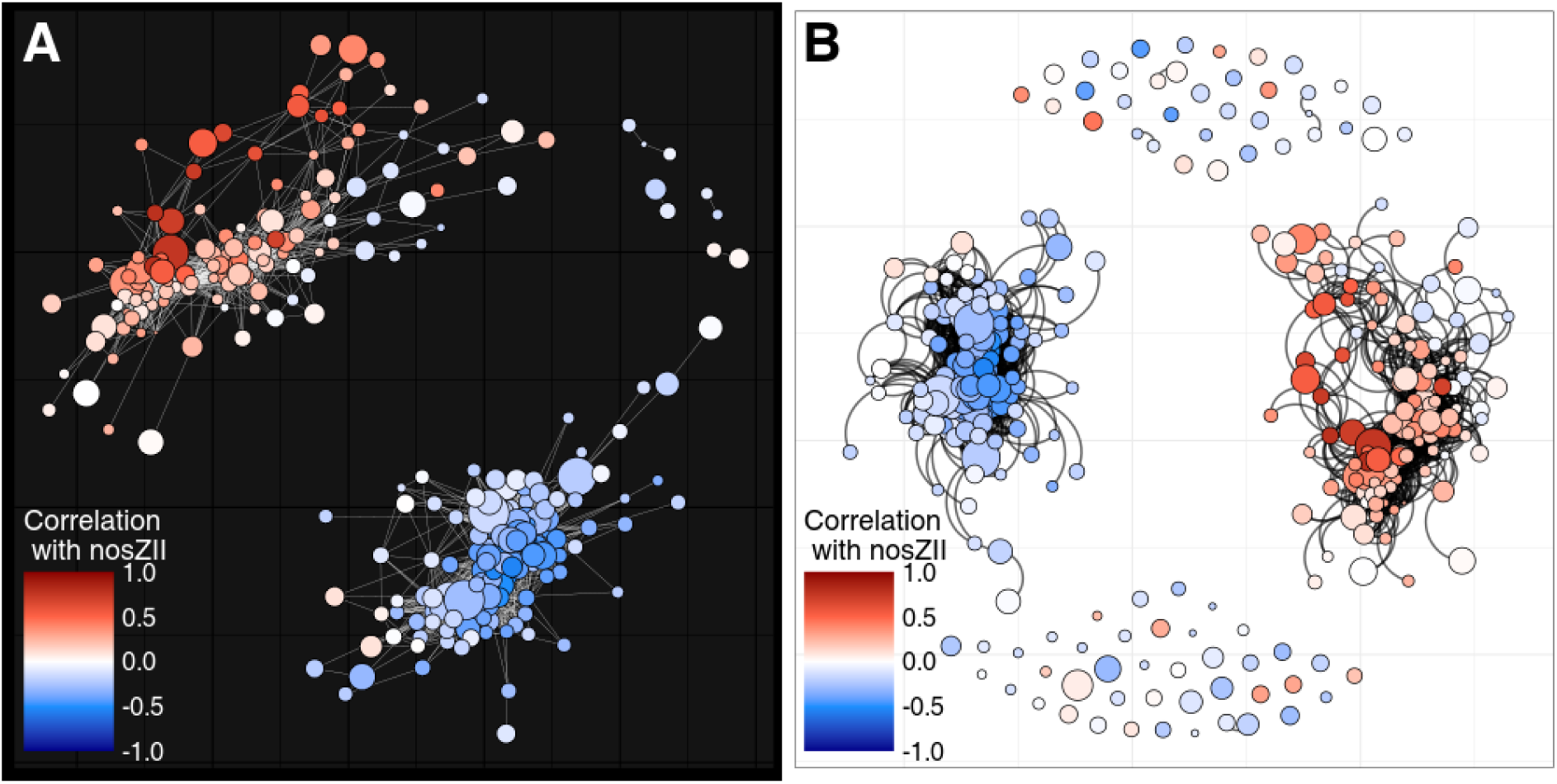
To illustrate customization options, we displayed the same correlation network with different aesthetic parameters (data from Brenzinger *et al*, same subnetwork as in Figure 4C-I). Color shows for each node (genus rank) the correlation between its relative abundance and the qPCR abundance of the *nosZII* gene. **A)** Network with default parameters (theme = “Dark”, edge strength = 0, edge alpha = 1, edge width = 0.1, random seed = 37, not showing isolated nodes). **B)** Same network with custom aesthetic parameters (theme = “Classic”, edge strength = 0.8, edge alpha = 0.6, edge width = 0.7, random seed = 16, showing isolated nodes).

**Figure S4:**
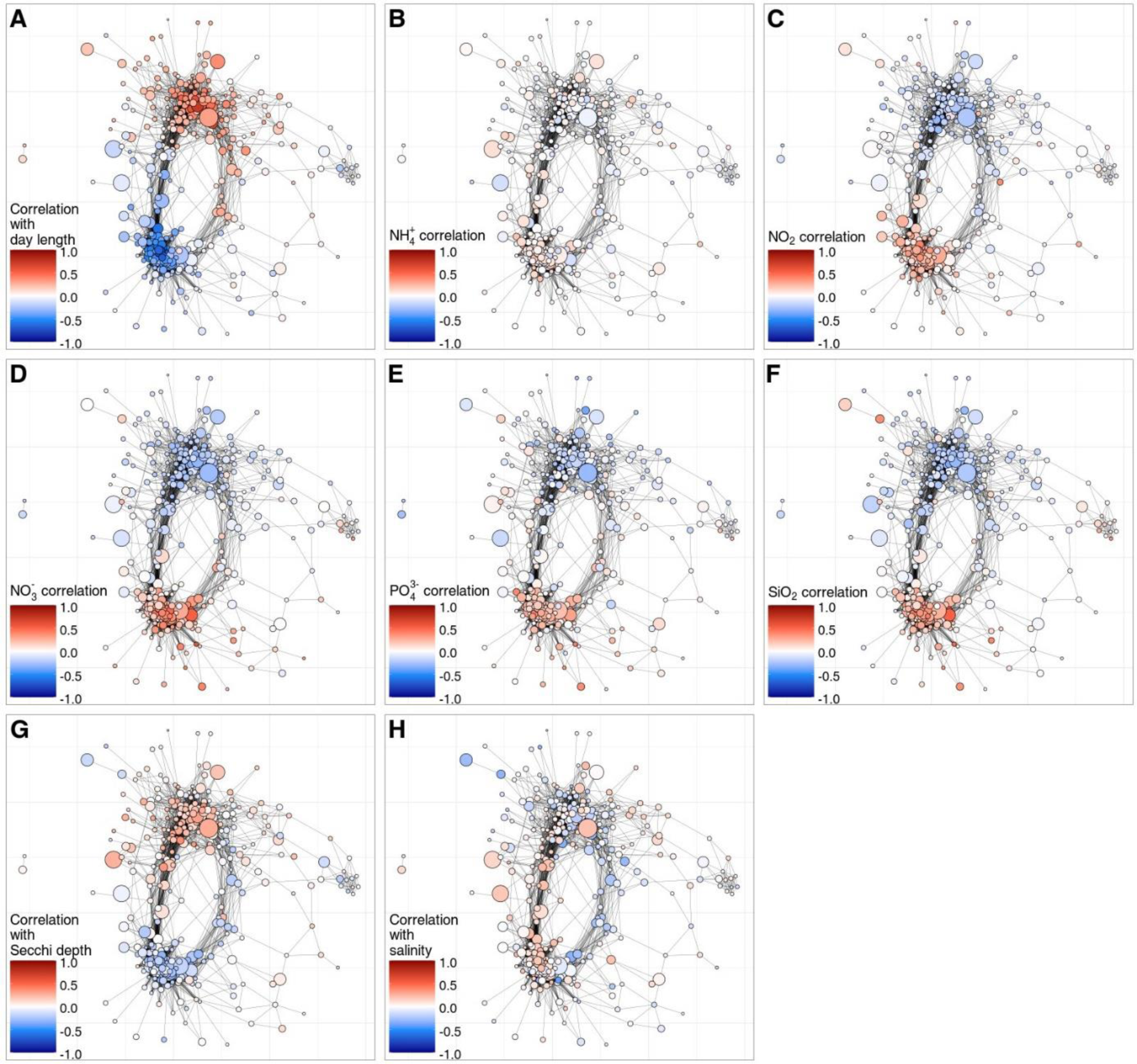
Inter-kingdom associations in a 10-year longitudinal study in the north-western Mediterranean sea (Deutschmann et al., 2023). Abundance table made by merging 16S and 18S table. Same network as in Figure 2C-F. Nodes are genera and edges are Spearman correlations > 0.4. Networks are colored by correlation between relative abundance of genera and: **A)** Hours of daylight **B)** 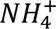 levels **C)** *NO*_2_ levels **D)** 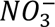 levels **E)** 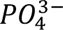 levels **F)** *SIO*_2_ levels **G)** Secchi depth (measure of water turbidity). **H)** salinity.

**Figure S5:**
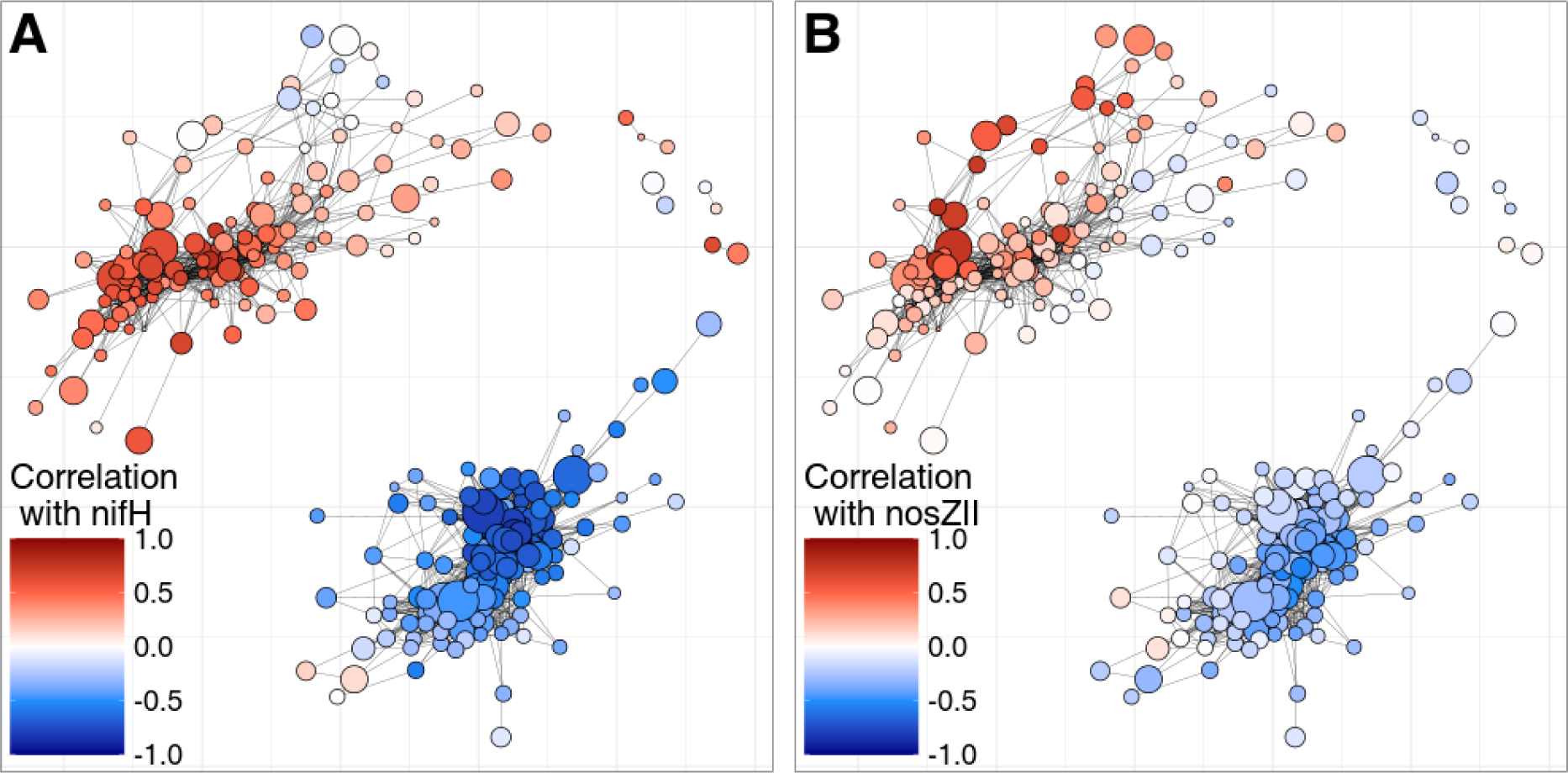
Correlations between genera and functional gene abundance for clay samples from Brenzinger *et al*. **A)** Network colored by Pearson correlation between genera and the qPCR abundance of the *nifH* gene. **B)** Network colored by Pearson correlation between genera and the qPCR abundance of the *nosZII* gene.

**Figure S6:**
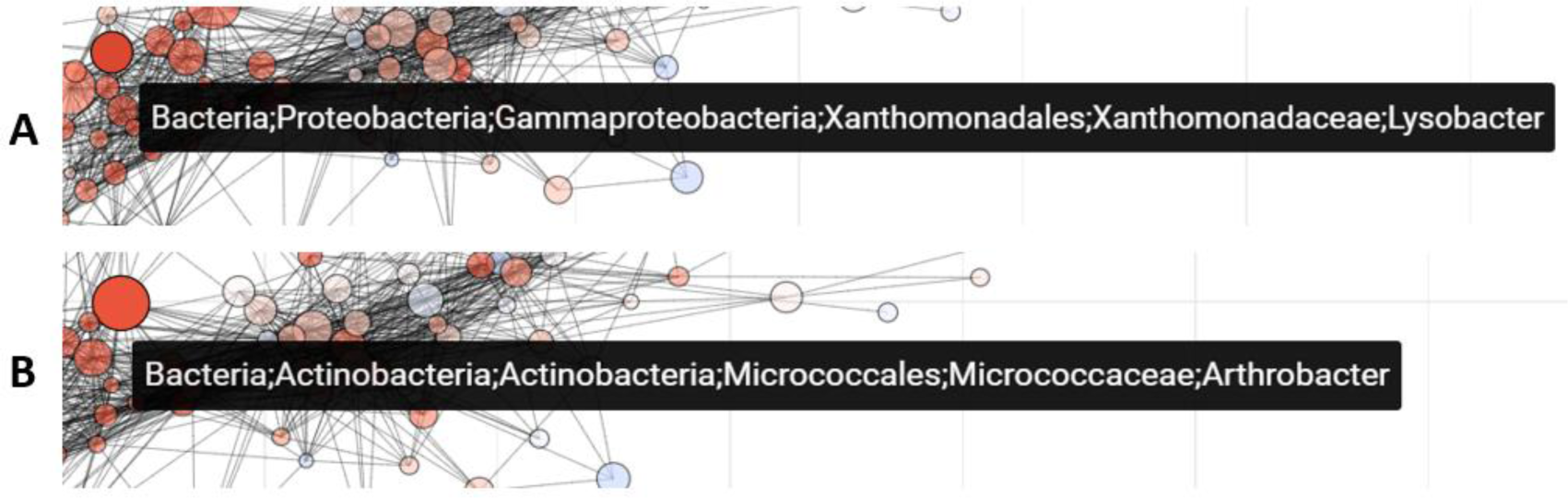
Mousing over the subnetwork with clay samples from Brenzinger *et al* reveals known plant-beneficial bacteria correlated with plant height. Color indicates correlation between the relative abundance of genera and the height of plants grown on the same plot. **A)** Mousing over a genus strongly correlated with plant height shows the taxonomy (genus *Lysobacter*). **B)** Mousing over a genus strongly correlated with plant height shows the taxonomy (genus *Achromobacter*).

**Figure S7:**
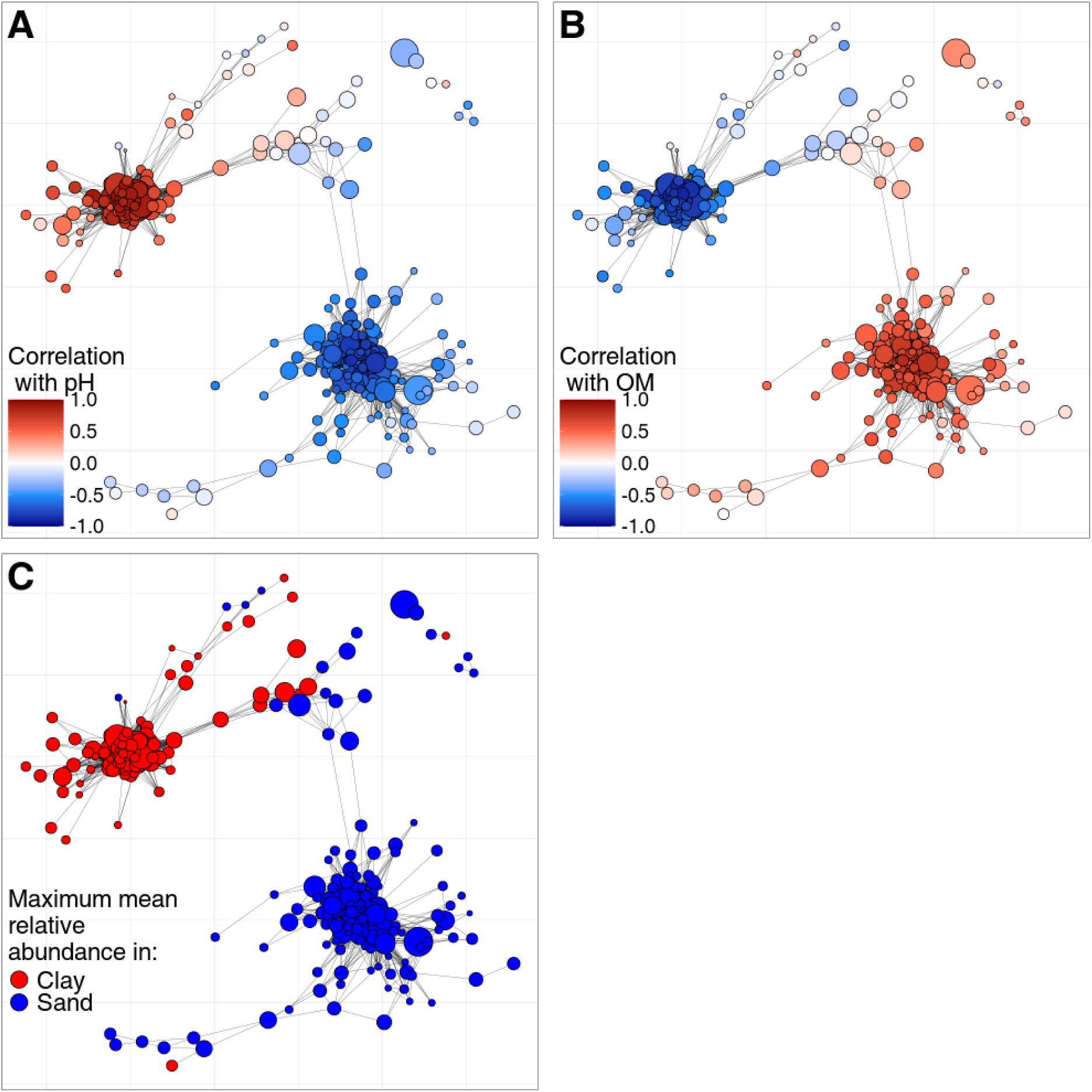
Correlated/linked environmental data can lead to visually convincing patterns but a causal effect on the microbiome cannot be concluded. Same network as in Figure 4A-B**. A)** Network colored by correlation of genus relative abundance with pH. **B)** Network colored by correlation of genus relative abundance with organic matter (OM). **C)** Network colored by preferred soil type of genera. A researcher seeing only panel A might be convinced the pH is structuring the microbiome. However, the pH in this dataset is strongly dependent on the soil type (mean and sd of pH in sand samples is 5.646, 0.183, mean and sd of pH in clay samples is 7.559, 0.084). Furthermore, the pH has a strong inverse correlation with the organic matter content of the soil (Pearson correlation = -0.91). Hence, whether and to what extent the pH, organic matter content, particle size of the soil or a combination thereof caused the observed microbiome structure cannot be determined from these networks.

