## Supplemental text S1: SpeSpeNet manual for "SpeSpeNet: An interactive and user-friendly tool to create and explore microbial correlation networks"

SpeSpeNet 1.0 manual

### Abstract

Correlation networks are commonly used to explore microbiome data. In these networks, nodes are taxa and edges represent correlations between their abundance patterns across samples. As clusters of correlating taxa (co-abundance clusters) often indicate a shared response to environmental drivers, network visualization contributes to system understanding. Currently, most tools for creating and visualizing co-abundance networks from microbiome data either require the researcher to have coding skills, or they are not user-friendly, with high time expenditure and limited customizability. Furthermore, existing tools lack focus on the relationship between environmental drivers and the structure of the microbiome, even though many edges in correlation networks can be understood through a shared relationship of two taxa with the environment. For these reasons we developed SpeSpeNet (Species-Species Network, https://tbb.bio.uu.nl/SpeSpeNet), a practical and user-friendly R-shiny tool to construct and visualize correlation networks from taxonomic abundance tables. The details of data preprocessing, network construction, and visualization are automated, require no programming ability for the web version, and are highly customizable, including associations with user-provided environmental data. This document outlines the functionalities of the SpeSpeNet tool.

### Dependencies

Using the web-version of SpeSpeNet (https://tbb.bio.uu.nl/SpeSpeNet) does not require any installation or programming. For the development version (available on github) the following libraries need to be installed:

1. Shiny
2. shinythemes
3. shinycssloaders
4. tidyverse
5. tidygraph
6. igraph
7. ggraph
8. ggdark
9. RColorBrewer
10. scales
11. plotly
12. phyloseq
13. shinyBS
14. MGnifyR
15. tools
16. pals
17. ggiraph
18. htmlwidgets
19. reshape2

Running the Launch_tool.R script will launch SpeSpeNet. Note that running SpeSpeNet requires an old (no longer available) installation of the MGnifyR package. In future we intend to develop a version that does not require the MGnifyR package. Until then it is strongly recommended to use the web version.

### 1. Navigating SpeSpeNet

SpeSpeNet has three tabs that can be navigated with the navigation bar (panels 1,2 and 3 in **Figure 1**). The “Network” tab is where data is uploaded and the network visualization is shown. The “Summary” tab shows the taxonomic composition and the relationship between co-response clusters/environmental categories and environmental metadata. The user can download .txt files of the network data in the “Raw data” tab.

### 2. Input data format

When starting SpeSpeNet the network tab is shown by default. Here, the user can input data, either as phyloseq objects (in a .rds file) or as three separate .txt files that contain the abundance table, the taxonomy table, and the environmental table (menu 4 in **Figure 1**). The data can be input as raw read counts, percentages, or fractions (menu 5 in **Figure 1**).

The .txt files need to be formatted in a standardized way: They should be **tab-separated**, the abundance table should have operational taxonomic units as rows and sample identifiers as columns, and the environmental table should have samples as rows and variables as columns. **The row names (containing OTU/ASV IDs) should match between the abundance table and the taxonomy table and be in the same order. The column names of the abundance table (containing sample IDs) should match the row names in the environmental data and be in the same order.** The column names of the taxonomic data should be taxonomic ranks (such as phylum, class, order, family, genus and species). To write row names and column names to a tab delimited .txt file, the following R code can be used:

write.table(matrix, "file.txt", sep = "\t", col.names = T ,row.names=T, quote = T).

Phyloseq objects must adhere to the same standards as the .txt files. SpeSpeNet can also download studies from the MGnify database if provided with a MGnify study accession number. The number of samples that will be downloaded from MGnify can be limited (menu 7 in **Figure 2**) in case downloading the whole study takes too long.

The user can choose to aggregate the OTUs/ASVs at the genus level (menu 6 in **Figures 1 and 2**) during upload. This is recommended for 16S datasets from habitats with high diversity like soil to constrain the number of nodes.

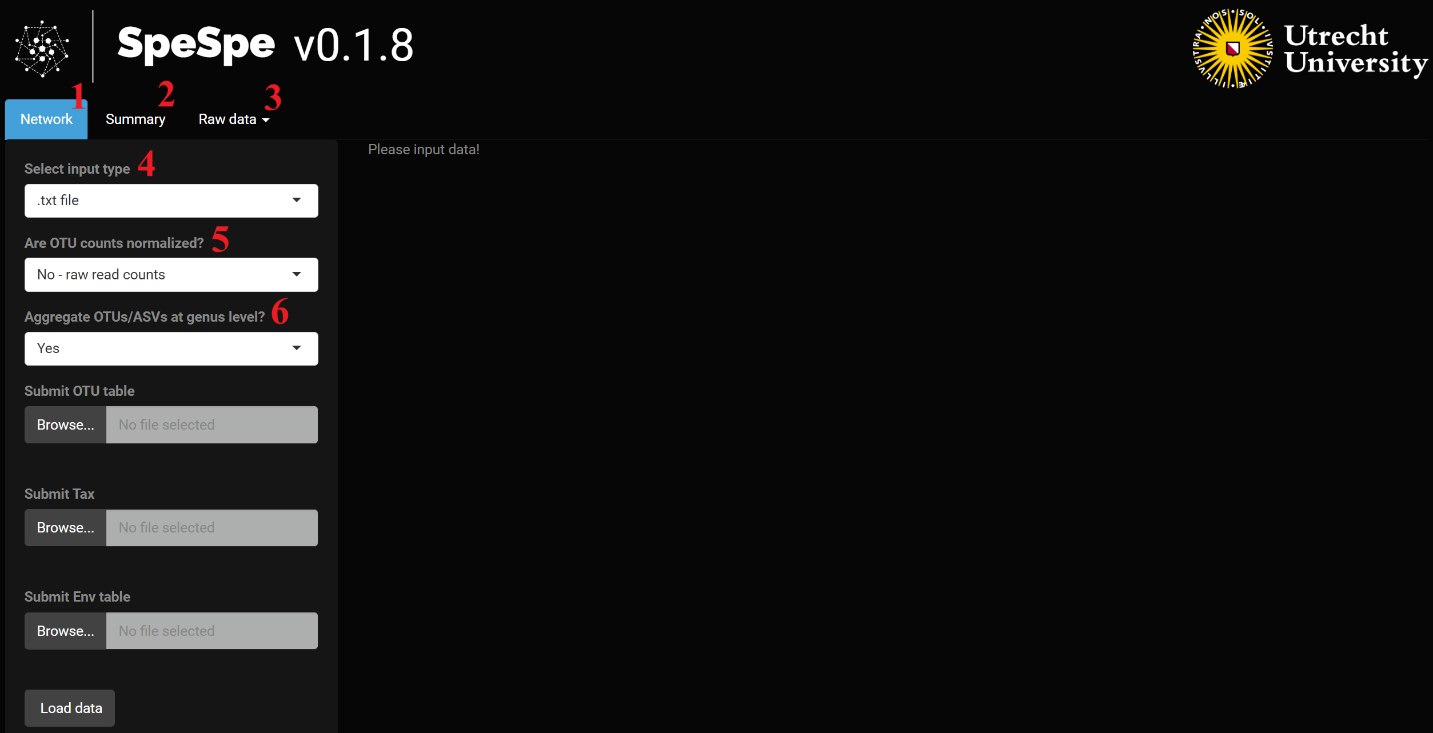

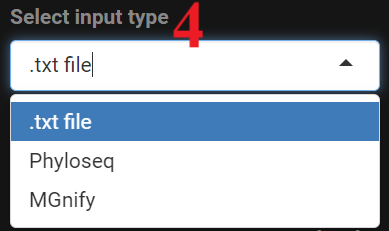

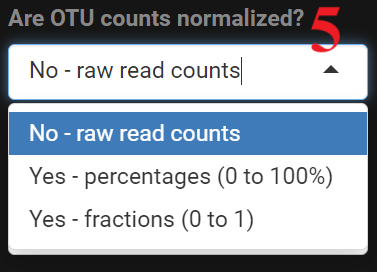

**Figure 1**. SpeSpeNet input options for .txt files

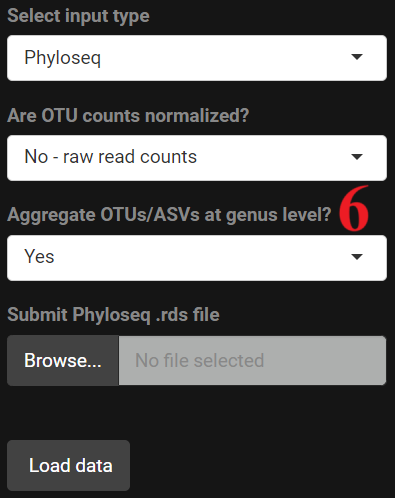

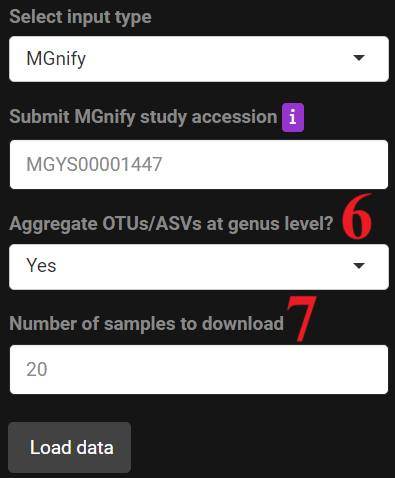

**Figure 2**. Input options for phyloseq objects and MGnify study accession numbers.

### 3. Network generation

After the data upload is finished SpeSpeNet constructs a co-abundance network using default settings (**Figure 3**). These settings for data filtering, network construction or network aesthetics are easily altered using buttons and sliders. Any change in the settings is instantly reflected in the network visualizations. Options for filtering include a threshold on the minimum number of samples a taxon should occur in (have non-zero read counts) to be included in the network (box 2 in **Figure 4**), and a threshold on the minimum relative abundance a taxon should have in at least one of the samples (box 3 in **Figure 4**). After filtering, the correlation matrix between the relative abundances of the taxa is calculated and used for network construction. The correlation method (Pearson, Spearman or Kendall) and correlation cut-off that defines edges can be chosen by the user (menu 4 in **Figure 4**). We recommend a rank-based correlation method (default is Spearman) due to the compositional nature of amplicon data.

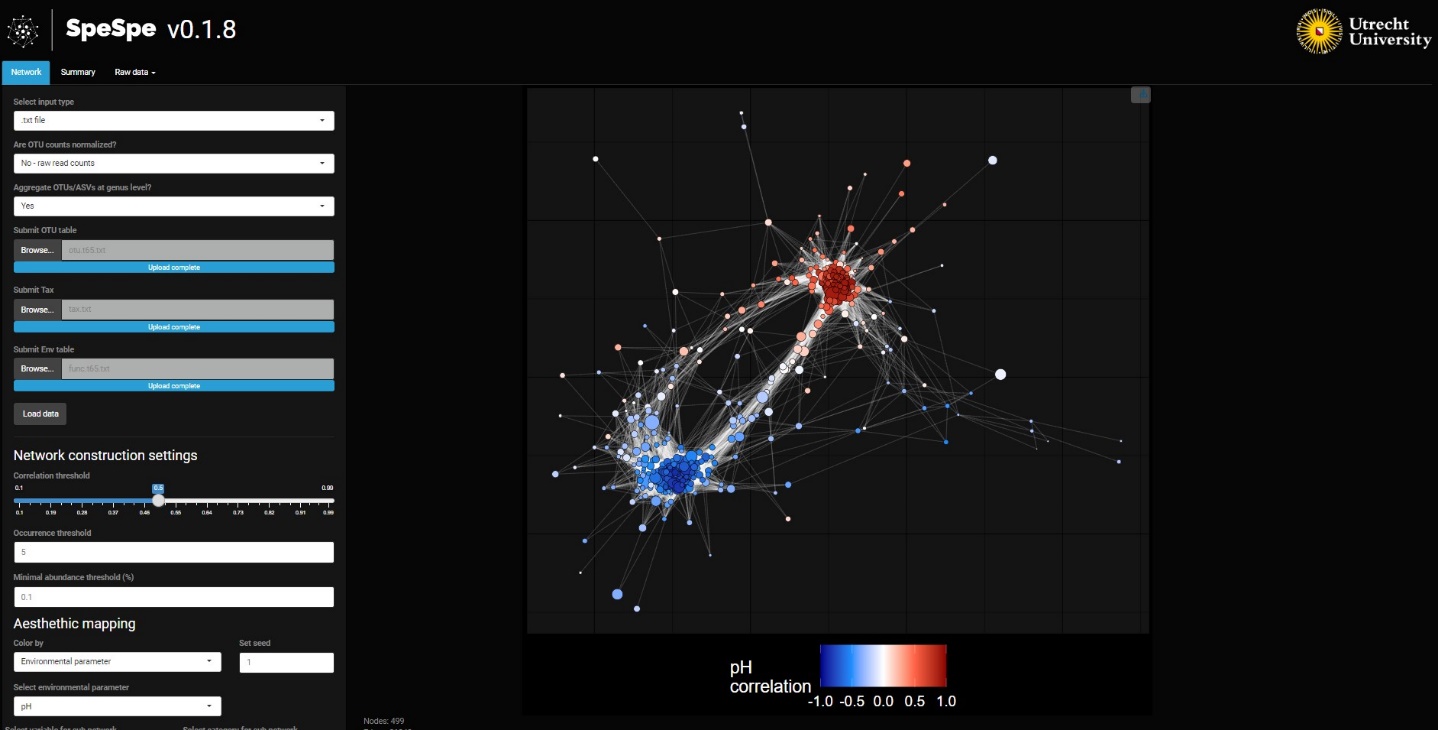

**Figure 3**. Network based on the default settings (data from Brenzinger *et al*, 2021).

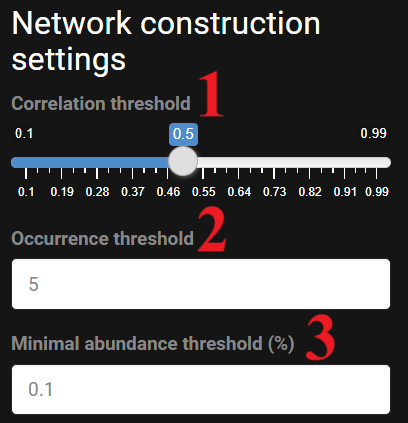

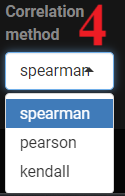

**Figure 4**. Network construction settings and correlation method options.

### 3. Coloring and changing aesthetics of the network

The network can be colored with three different kinds of variables: 1) k-means clustering, 2) taxonomy and 3) relationship with an environmental variable (menu 1 in **Figure 5**). In case 1) the k-means algorithm is applied on the correlation matrix of dimensions taxa x taxa. The user can specify the number of clusters made by the k-means algorithm (box 2 in **Figure 5**). The network is then colored according to the clustering. In case 2) the network can be colored on taxonomy at a chosen taxonomic rank (menu 3 in **Figure 5**). SpeSpeNet will show the fifteen most abundant clades and lump the rest together as “Other” to constrain the size of the legend. Finally, in case 3) the network is colored on an environmental variable in the metadata (menu 2 in **Figure 5**). If a numeric environmental variable is selected (e.g. pH), the color of the nodes represents the correlation of the relative abundance of the taxa to the environmental variable. If a categorical variable is selected (e.g. healthy/diseased or sand/clay), SpeSpeNet will color on the category in which the taxon has the highest mean relative abundance. The network is interactive and shows the taxonomy when hovering the cursor over a node (regardless of the coloring of the network) (**Figure 6**).

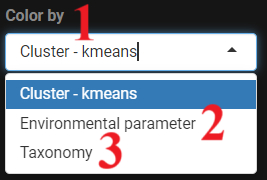

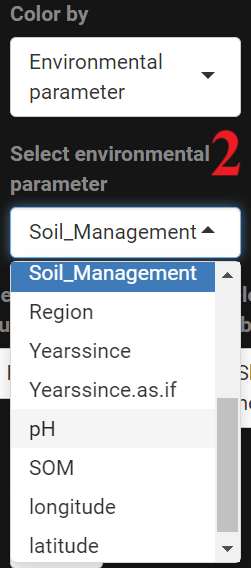

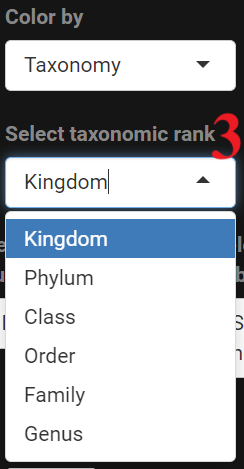

**Figure 5.** Options to color the network.

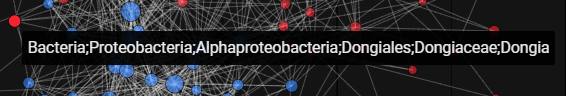

**Figure 6.** Hovering the cursor over a node shows the taxonomic classification.

Options to change the coloring or aesthetics of the network (**Figure 7**) are found underneath the network construction options. Aesthetic options include the background theme, curvedness of edges, edge width, edge transparency, and whether isolated nodes (nodes without edges) should be plotted (boxes 6-10 in **Figure 7**). The random seed (box 3, **Figure 7**) changes the orientation of the network but will not affect the nodes, edges or clustering of the network.

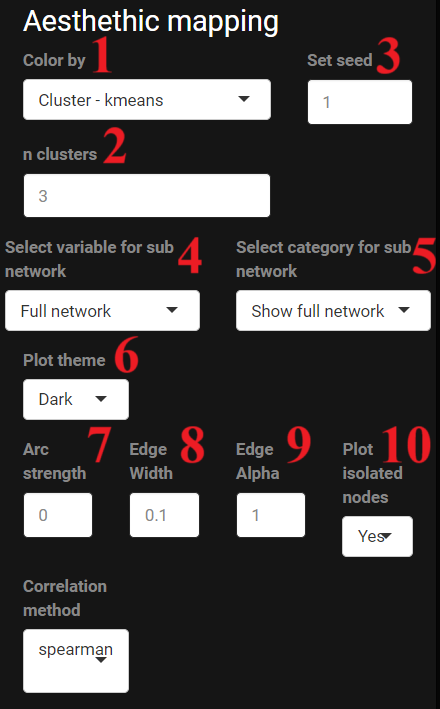

**Figure 7.** SpeSpeNet aesthetic mapping options.

If there are categorical environmental variables, the user can zoom in on a specific category (menu 4 in **Figure 7**). This is only possible for categories with at least eight samples. A new correlation matrix and network is then constructed using only the samples in the chosen category. As an example, we show a network constructed using the entire dataset and a network constructed using clay samples only (data from Brenzinger *et al*, 2021) (**Figure 8**).

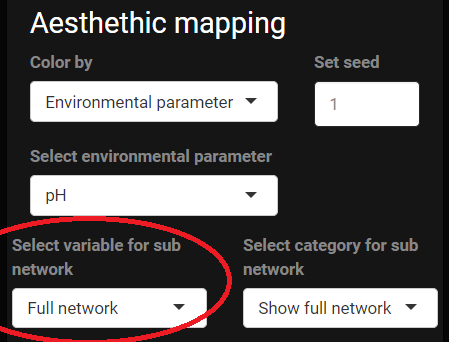

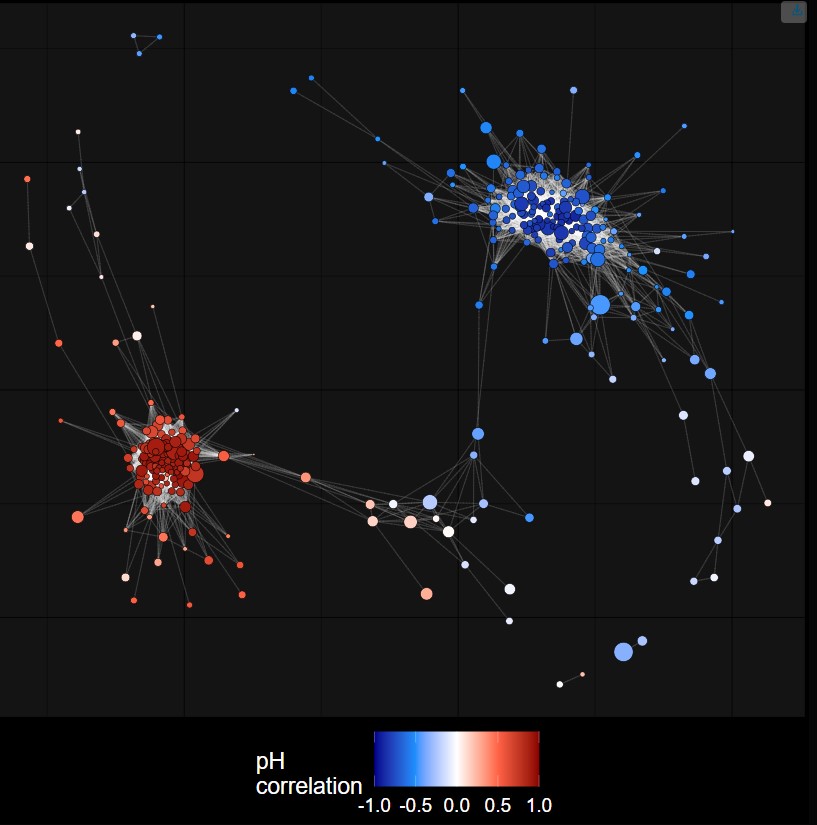

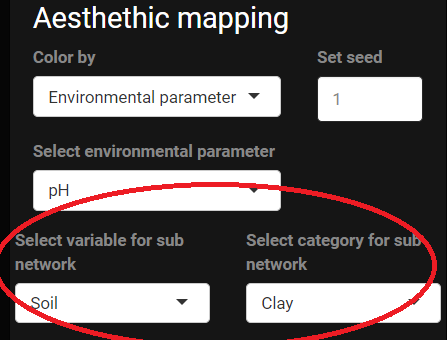

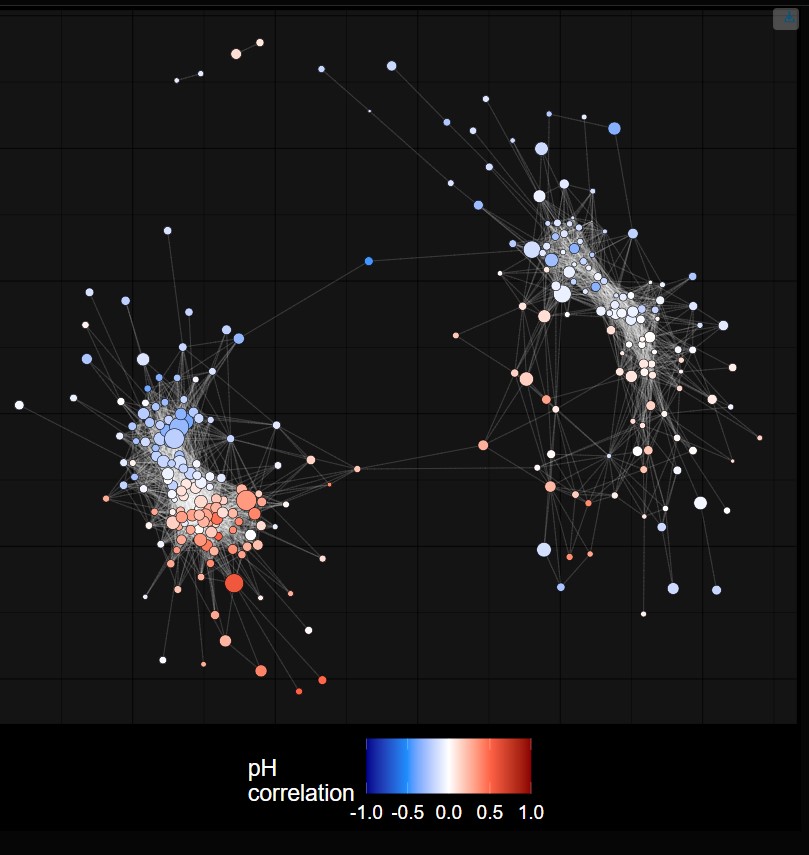

**Figure 8**. Network made using the full dataset and a subnetwork constructed using only samples from clay soils (data from Brenzinger *et al*, 2021).

### 4. Downloading the network visualization

The network can be downloaded as a .png with the button in the top right corner (small button with blue arrow in networks in **Figure 8**).

### 5. The summary tab

The summary tab contains a barplot of the taxonomic composition and a boxplot of the Pearson correlations between the relative abundance of genera and a user-specified continuous environmental variable (**Figure 9**).

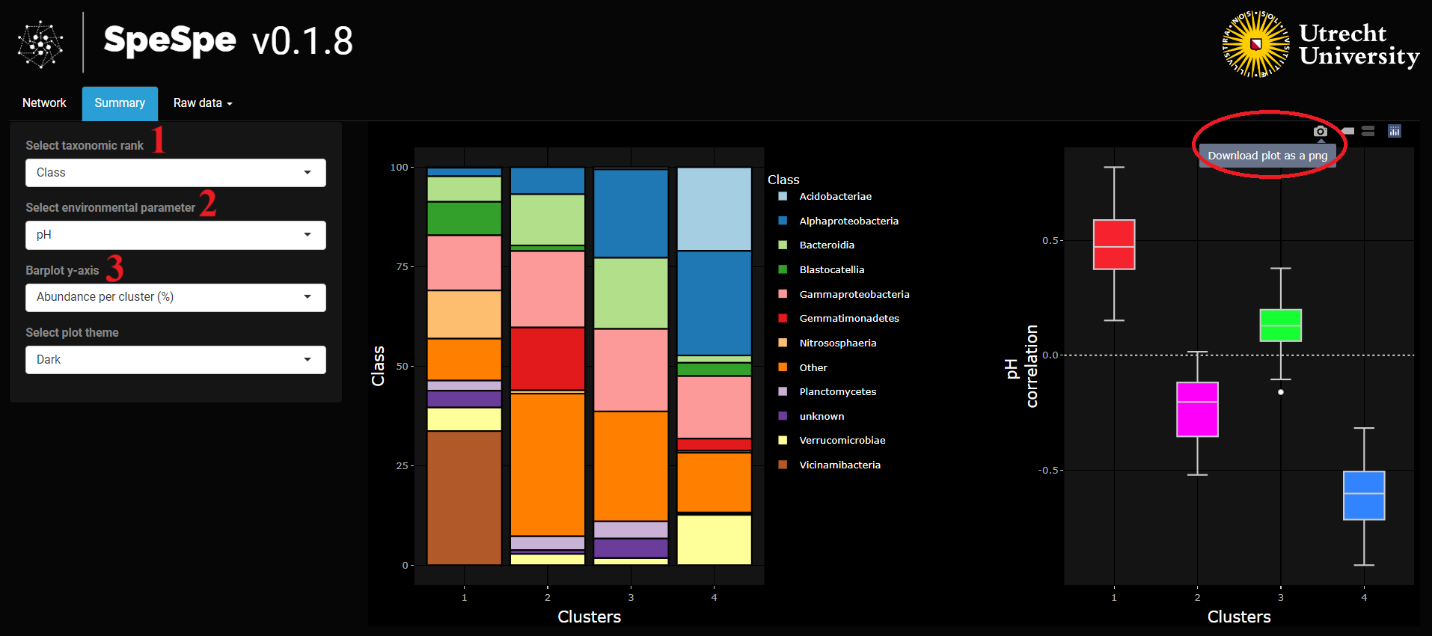
**Figure 9.** Overview of the Summary tab.

If the network is colored on k-means clustering or a categorical environmental variable, the barplot shows the taxonomic composition of each individual cluster or category (increasing or decreasing the number of clusters in the network panel has an immediate effect on the summary tab too). If the network is colored on taxonomy or a numerical environmental variable, the barplot in the summary tab shows the overall taxonomic composition of the entire dataset. The user can choose at which taxonomic rank they want the composition plotted (menu 1 in **Figure 9**) and (if using k-means clusters) whether the y-axis shows the mean relative abundance over all samples or per cluster (menu 3 in **Figure 9**) (Compare top panel with bottom panel in **Figure 10**). The barplot shows the mean relative abundance as a percentage when hovering the mouse over the bar plot (**Figure 10**).

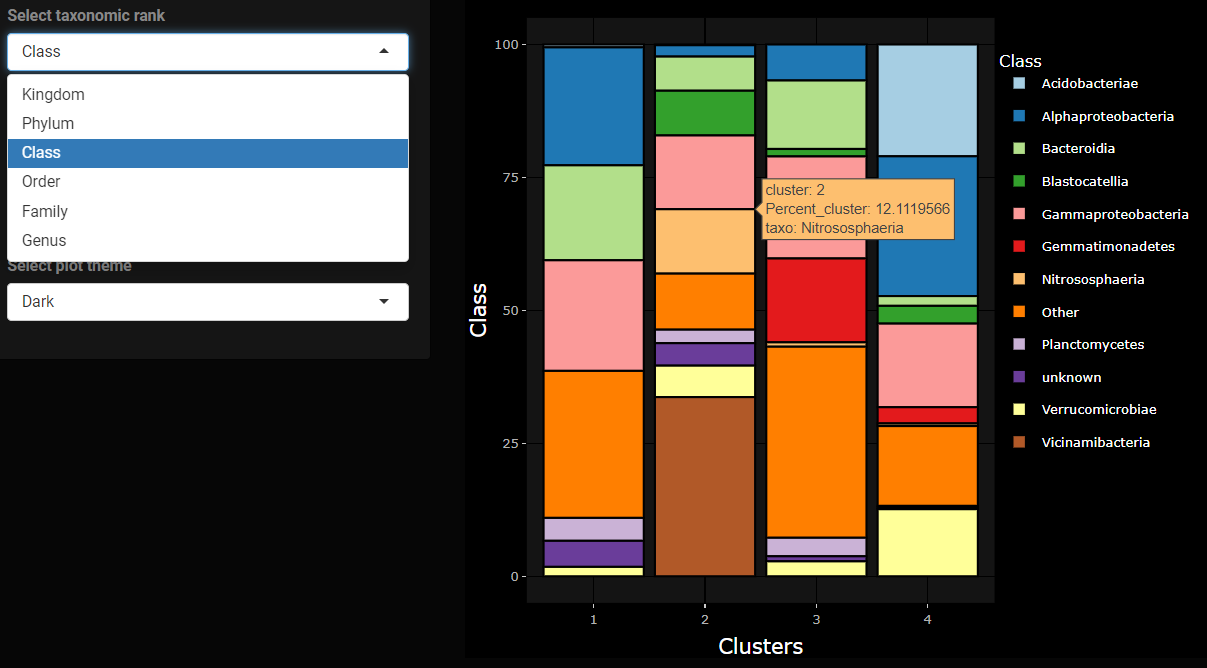

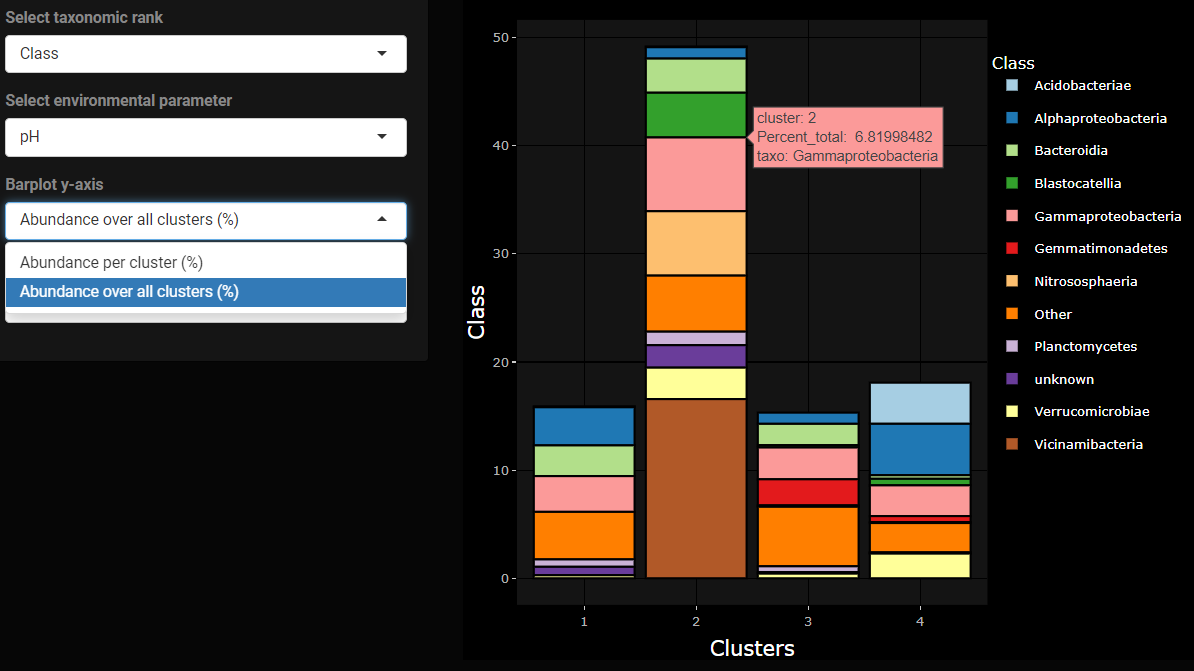

**Figure 10.** The y-axis options and interactivity of the barplot in the summary tab.

The boxplot shows the correlations of the relative abundance of the taxa (nodes in the network) per cluster or over the entire dataset with a numeric environmental variable in the environmental metadata (menu 2 in **Figure 9**). Hovering the mouse over a box shows summary statistics of the distribution of the correlations (minimum and maximum value, median and the two remaining quartiles). The barplot and boxplot can be downloaded together as a .png with the button in the top right (circled in red in **Figure 9**).

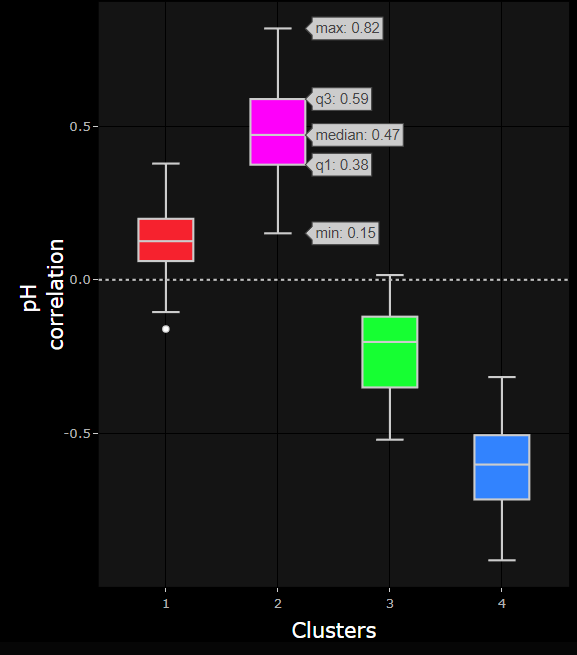

**Figure 11.** The interactivity of the boxplot.

### 5. The raw data tab

The third tab is the “Raw data” tab, from which network data can be previewed and downloaded. The user can download the correlation matrix, node data, and edge data in the form of .txt files.

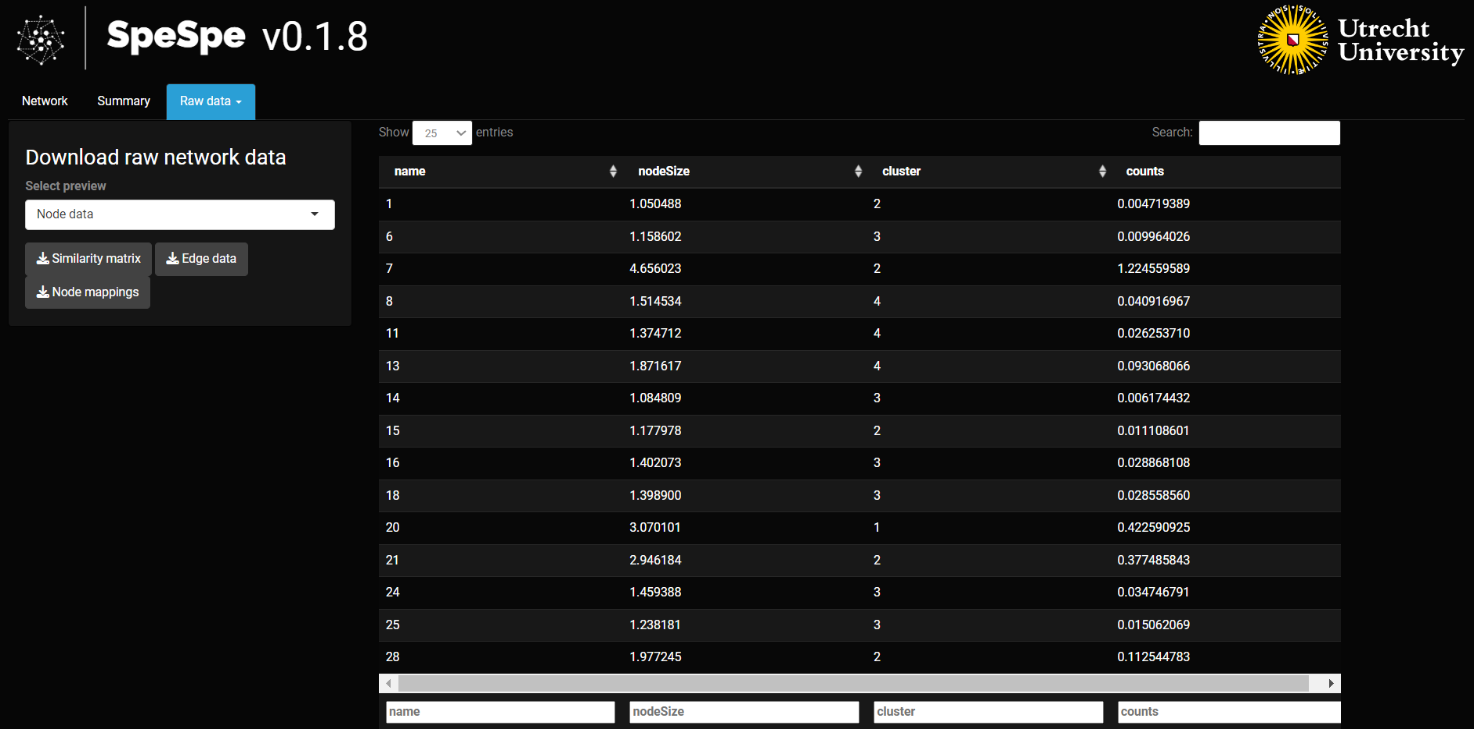
**Figure 12.** Overview of the Raw data tab.
